# Reproductive history determines *ErbB2* locus amplification, WNT signalling and tumour phenotype in a murine breast cancer model

**DOI:** 10.1101/2020.10.30.361998

**Authors:** Liliana D Ordonez, Lorenzo Melchor, Kirsty R. Greenow, Howard Kendrick, Giusy Tornillo, James Bradford, Peter Giles, Matthew J. Smalley

## Abstract

Understanding the mechanisms underlying tumour heterogeneity is key to development of treatments that can target specific tumour subtypes. We have previously targeted CRE recombinase-dependent conditional deletion of the tumour suppressor genes *Brca1*, *Brca2*, *p53* and/or *Pten* to basal or luminal ER− cells of the mouse mammary epithelium. We demonstrated that both the cell-of-origin and the tumour-initiating genetic lesions co-operate to influence mammary tumour phenotype. Here, we use a CRE-activated HER2 orthologue to specifically target HER2/ERBB2 oncogenic activity to basal or luminal ER− mammary epithelial cells and carry out a detailed analysis of the tumours which develop. We find that in contrast to our previous studies, basal epithelial cells are refractory to transformation by the activated *NeuKI* allele, with mammary epithelial tumour formation largely confined to luminal ER− cells. Histologically, the majority of tumours that developed were classified as either adenocarcinomas of no special type or metaplastic adenosquamous tumours. Remarkably, the former were more strongly associated with virgin animals and were typically characterised by amplification of the *NeuNT/ErbB2* locus and activation of non-canonical WNT signalling. In contrast, tumours characterised by squamous metaplasia were associated with animals that had been through at least one pregnancy and typically had lower levels of *NeuNT/ErbB2* locus amplification but had activated canonical WNT signalling. Squamous changes in these tumours were associated with activation of the Epidermal Differentiation Cluster. Thus, in this model of HER2 breast cancer, cell-of-origin, reproductive history, *NeuNT/ErbB2* locus amplification, and the activation of specific branches of the WNT signalling pathway all interact to drive inter-tumour heterogeneity.

## Introduction

The ERBB2/HER2 receptor tyrosine kinase (a member of the epidermal growth factor receptor family) is amplified and overexpressed in 20-30% of human breast cancers, leading to an aggressive form of the disease [1]. Therapeutic strategies which block HER2 activity, either with antibodies targeted to the extracellular domain or small molecules blocking intracellular kinase activity, have substantial benefit [2] although primary or acquired resistance remains an issue [3]. HER2 activity has been associated with stem cell-like behaviour in normal breast and breast cancer [4, 5] but the cell-of-origin of HER2 breast cancers remains unclear.

Understanding how cell-of-origin and genetic lesions interact to drive tumour behaviour and heterogeneity is key to development of treatments which can be targeted to specific tumour subtypes. In the mammary epithelium, potential cells of tumour origin include basal cells, luminal oestrogen receptor negative (ER-) progenitors and luminal ER+ cells (largely differentiated hormone sensing cells although including a small proportion of progenitors) [6]. In previous studies of mammary tumour origin, we have used genetically engineered mouse models in which CRE recombinase-dependent conditional deletion of the tumour suppressor genes *Brca1*, *Brca2*, *p53* and/or *Pten* were targeted to basal or luminal ER− cells using the *Krt14* or *Blg* promoters, respectively. We demonstrated that both the cell-of-origin and the tumour-initiating genetic lesions co-operate to influence the tumour behaviour, with a more diverse range of tumour phenotypes arising from luminal ER− cells but basal-origin tumours having significantly shorter latency [7, 8].

Here, we have addressed whether cell-of-origin similarly affects the development and phenotype of *HER2*-amplified tumours. We have taken advantage of the *NeuKI* allele, in which an activated mutant variant of *Neu* (*NeuNT*), the rat *Erbb2/Her2* orthologue, has been knocked into the endogenous *Erbb2* locus. *NeuNT* is therefore expressed under the control of the endogenous promoter but only when an upstream *loxP-*flanked neomycin cassette is excised by CRE recombinase activity [9]. Using this allele, we have been able to target NeuNT activity to basal or luminal ER− cells using our established *Krt14Cre* and *BlgCre* lines. We have carried out a detailed analysis of *Krt14Cre-NeuKI* and *BlgCre-NeuKI* mice, assessing tumours from virgin and parous animals. For comparison, we have used both normal non-transgenic tissue and tumours from the *MMTV-Neu* model, in which *Neu* expression is driven constitutively by a very strong mammary promoter [10].

We find that in contrast to our previous studies, basal epithelial cells are refractory to transformation by the activated *NeuKI* allele, with mammary epithelial tumour formation largely confined to luminal ER− cells. Most tumours arising from the latter population were classified as either adenocarcinomas of no special type (AC(NST)) or metaplastic adenosquamous carcinomas (ASQC). Remarkably, the proportion of ASQCs arising from this cell type depended on the reproductive history of the animal, with ASQC tumours strongly associated with animals that had been through at least one pregnancy. Furthermore, while AC(NST) tumours were typically characterised by high amplification of the *NeuNT/ErbB2* locus and activation of non-canonical WNT signalling, ASQC tumours typically had lower levels of *NeuNT/ErbB2* locus amplification but activated canonical WNT signalling.

Thus, reproductive history affects *NeuNT/ErbB2* locus amplification and the activation of specific branches of the WNT signalling pathway and ultimately drives inter-tumour heterogeneity in this murine model of human HER2 breast cancer. Importantly, our findings also show that some cell types are intrinsically resistant to transformation by particular oncogenic drivers, suggesting at least one mechanism behind why certain mutations only cause cancer in a distinct range of target organs.

## Methods

### Experimental mice

All procedures were conducted according to UK Home Office regulations and ARRIVE guidelines and under the authority of appropriate licences following local ethical review by the Cardiff University Animal Welfare Ethical Review Body. Mice carrying the targeted *NeuKI* allele (see schematic in **Figure S1**) were kindly supplied by Professor Margaret Frame (University of Edinburgh) with permission from the originator (Professor Bill Muller, McGill University, Montreal). Mice of two genotypes, *Krt14Cre-NeuKI* and *BlgCre-NeuKI* were bred and maintained on a mixed FVB/C57Bl6 background. Some animals from each genotype were aged as virgin animals, others went through one or more pregnancies (full details of all animals used are provided in **Table S1**). *Cre* and *NeuKI* alleles were maintained as heterozygous loci; the non-recombined *NeuKI* locus is inactive and homozygosity of this allele is lethal. Genotyping primers and conditions are given in **Table S2**.

The well-established *MMTV-NeuNDL* mouse line (a kind gift of Don White) [10] was maintained to enable comparison with a widely used model of *HER2* disease driven purely by strong over-expression. For analysis of gene expression in wild-type mice, C57Bl6 animals were used.

### Histology and Immunohistochemistry

Tumour tissue was fixed in ice cold 4% neutral buffered formalin for 24 hours before being processed into paraffin blocks according to standard procedures. Tissue sections (5μm) were either stained using haematoxylin & eosin (H&E) for histological analysis, or were used for immunohistochemistry as described [7]. The following antibodies were used: anti-K14 (1:500; Abcam), anti-K18 (1/5; Progen Biotechnik), anti-ΔNp63 (1:100; Abcam), anti-ERα (1:500; Vector Labs), anti-PRA (1:500; Thermo Scientific), anti-PRB (1:75; Abcam), anti-βcatenin (1:200; Becton Dickinson), anti-ERBB2 (1/500; Calbiochem). Mouse tumour phenotyping based on H&E appearance and levels and staining patterns of ΔNp63, keratins, ER and PR was carried out as we have previously described [7, 8] according to the WHO classification of tumours in the breast [11] by MJS with support and advice from Professor Barry Gusterson FRCPath.

### Quantitative real-time RT-PCR

Total RNA was isolated from tissues using Trizol (Life Technologies, Paisley, UK) or the RNeasy kit (QIAGEN, Manchester, UK) according to manufacturer’s protocol. One microgram of RNA was reverse transcribed using Quantitect (Life Technologies Ltd., Paisley UK) according to the manufacturer’s protocol. Gene expression analysis was carried out using TaqMan Universal PCR mastermix according to the manufacturer’s protocol (Applied Biosystems, Life Technologies Ltd., UK). TaqMan probes (**Table S2**) were obtained from Taqman gene expression assays (Applied Biosystems). Data analysis was carried out using QuantStudio 7 (Applied Biosystems). Relative expression levels of target genes were calculated using the ΔΔC_t_ method as described [12] with *Actb*, *B2m* and *Gapdh* as the endogenous controls.

### Digital droplet PCR (ddPCR) for Copy Number Variation (CNV)

DNA from tumour and organoid samples were extracted using the Gentra Puregene Tissue Kit (QIAGEN) as per manufacturer’s guidelines. Test samples were diluted to 300 ng/μl in RNase/DNase free water and CNV assays were prepared for partitioning by adding 300 ng of DNA per CNV assay probe (**Table S2**) to a restriction enzyme mix containing 0.01 U each of PacI, PsiI and MluI. Samples were incubated for 5 minutes at 37°C before chilling and addition of ddPCR buffer and reference probe. This mix was then aliquoted into PCR strips containing individual test probes and RNase/DNAse free water to a total of 25 μl per sample. The partition mix was then transferred into the appropriate wells of a BioRad (BioRad, Watford, Hertford, UK) droplet generation cartridge, followed by addition of droplet oil to respective wells, the cartridge gasket sealed and droplets generated using the BioRad QX200 Droplet Generator. Droplets were subsequently transferred to a PCR plate for amplification consisting of one initial cycle of 95°C for 10 minutes followed by 40 cycles of 94°C for 30 seconds and 60°C for 1 minute with a final denaturing step of 98°C for 10 minutes. Plates were then analysed within 1 hour of droplet generation on a BioRad QX200 droplet reader. Analysis was performed using the QuantaSoft Analysis program (BioRad).

### Next Generation Sequencing (NGS) analysis

Ten *BlgCre NeuKI* tumours with representative phenotypes were selected for RNA Seq-based analysis of gene expression. Genomic DNA was isolated from the same ten tumours, together with matched spleens, for exome sequencing. NGS analysis was carried out by the Institute of Cancer Research Tumour Profiling Unit (Chester Beatty Laboratories, Fulham Road, London, UK).

For RNASeq, genomic DNA was removed from 500ng of total RNA using the genomic DNA eliminator column from RNeasy Plus Micro Kit (QIAGEN) and rRNA removed using RiboZero (Epicentre, Wisconsin, USA) following manufacturer’s instructions. Strand-specific libraries were created using NEBNext Ultra Directional RNA Library Prep Kit for Illumina (NEB, Hitchin, Herts, UK) and 20ng of the rRNA depleted RNA.

For ExomeSeq, genomic DNA (200-600ng) was fragmented to 200bp using a Covaris E Series and the resultant libraries were subjected to DNA Capture using SureSelect XT Mouse All Exon kit (Agilent, Stockport, Cheshire) following manufacturer’s instructions.

Final libraries from both RNASeq and ExomeSeq preparations were quantified using qPCR and clustered at a molarity of 14.5 pM, sequencing was performed on an Illumina HiSeq 2500 using 2×76 cycles of version 3 SBS chemistry.

Raw and processed RNAseq and exome seq data have been deposited at GEO with the overall accession number GSE162348.

### Bioinformatic analysis of RNASeq data

Raw FASTQ sequence files were quality control checked using FastQC (http://www.bioinformatics.babraham.ac.uk/projects/fastqc). To estimate gene expression, reads were aligned to the mouse genome (build 38) using StarAlign [13] with no more than three mismatches, and only uniquely mapped reads permitted. Reads whose ratio of mismatches to mapped length was greater than 0.10 were also discarded. All other parameters were set to their defaults for un-stranded alignment. The expression level, based on Fragments Per Kilobase per Million fragments mapped (FPKM), of each gene present in the mouse GTF annotation (build 38) file downloaded from Ensembl [14] was estimated using Cufflinks [15] with library type defined as “fr-unstranded”. All other parameters were set to defaults. Read counts across each gene were calculated by HTSeq-count [16], and input to DESeq2 [17] to detect differential expression. FPKMs from multiple samples were merged to generate a gene-by-sample matrix using a custom Perl script for input into downstream signature scoring algorithms.

Non-negative matrix factorization (NMF) was used to cluster tumour gene expression data. Only the most highly expressed and variable genes were chosen for clustering according to the following criteria: (mean FPKM+SD)>1.00 and CV>0.10, where CV = coefficient of variation. The underlying principle of NMF is dimensionality reduction in which a small number of meta-genes, each defined as a positive linear combination of the genes in the expression data, are identified and then used to group samples into clusters based on the gene expression pattern of the samples as positive linear combinations of these meta-genes. Using the R package NMF [18], factorization rank k was chosen by computing the clustering for k = 2–6 against 50 random initializations of both the actual and a permuted gene expression matrix, and selecting the k value achieving the largest difference between cophenetic correlation coefficients calculated from the actual and permutated data. For further visual confirmation of a sensible choice of k, consensus matrices were generated corresponding to different k values. To achieve stability, the NMF algorithm was then run against 200 perturbations of each gene expression matrix at the chosen value of k = 3.

With the exception of sample MS1218.2, ESTIMATE analysis [19] of the RNAseq data demonstrated that the samples had better than >80% purity of tumour cells.

### Mutation calling in ExomeSeq data

Read pairs were mapped against the mouse genome (build 38) using BWA “mem” algorithm with default parameters [20]. The resulting bam files were then pre-processed in preparation for somatic mutation detection using the Genome Analysis Toolkit (GATK) v3.5 best practice pipeline [21] and dbsnp version 144 in the base recalibration step [22]. MuTect v1.1.7 was then applied to compare the resulting bam files from tumour and matched normal tissue to call somatic mutations [23]. Mutations were annotated using the Ensembl Variant Effect Predictor [24] using the canonical transcript, and non-silent protein coding mutations taken forward for further consideration.

### Copy number profiling using ExomeSeq data

Read pairs were mapped against the mouse genome (build 38) using BWA “mem” algorithm with default parameters [20]. Duplicate reads were removed, as were reads achieving mapping quality below 37. Depth of coverage at each position of all exons annotated according to the UCSC Genome Browser was calculated using GATK “DepthOfCoverage” tool [21], and the resulting tumour and normal profiles input to ExomeCNV R package using default parameters [25]. Gene level log2 copy number ratios were then parsed using custom Perl scripts, with gains achieving log2 ratio > 4.00, and losses < −2.50 taken forward for further consideration.

### Statistics

Statistical analysis was carried out in GraphPad Prism. Significance of survival curves was determined using Log Rank Mantel-Cox test and the Gehan-Breslow-Wilcoxon test. Statistics for differences between tumour numbers per animal used the Mann-Whitney test. Comparison of Neu amplification status used two-tailed t-test with Welch’s correction on Log10 transformed values. Significance of gene expression differences analysed by qrtPCR were determined from 95% confidence intervals [26].

## Results

### Luminal progenitors are sensitive and basal cells resistant to NeuKI allele-dependent tumourigenesis

*Krt14Cre* targets CRE expression/activity primarily to basal cells and *BlgCre* primarily to luminal ER negative cells in the mammary epithelium [8]. To test the contribution of cell-of-origin to tumour phenotype in HER2 mouse models, cohorts of *Krt14Cre-NeuKI*, *BlgCre-NeuKI* and *MMTV-NeuNDL* mice were aged, together with a control group carrying the *NeuKI* allele but no *Cre* transgene. A subset of the *BlgCre* and *Krt14Cre* cohorts underwent one or more pregnancies.

Full details of the animals in the cohorts are provided in **Table S1**. These included 33 *MMTV-NeuNDL* mice (26 virgin, 7 parous); 42 *BlgCre-NeuKI* mice (23 virgin, 19 parous); 27 *K14Cre-NeuKI* mice (18 virgin, 9 parous); and 12 control mice carrying the *NeuKI* allele but no *Cre* transgene (all virgin). Kaplan-Meier curves for overall survival are shown in **Figure 1A** and statistical comparisons are provided in **Table S3**. The penetrance of the different phenotypes within each line is shown in **Figure 1B**.

**Figure 1:**
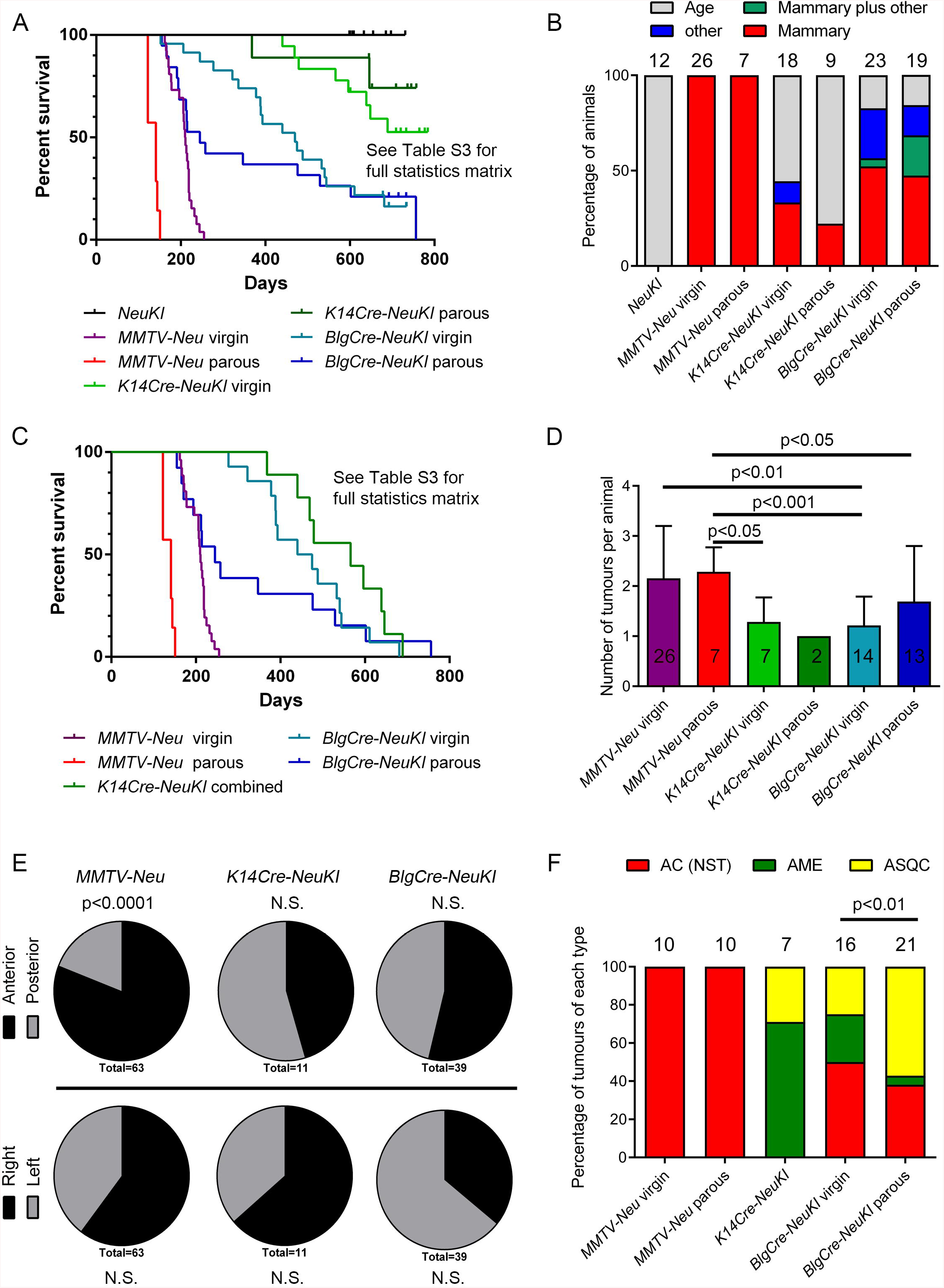
Features of tumour cohorts. **(A)** Kaplan-Meier survival curves (time to euthanasia for any reason) for all cohorts. Statistical significances between the cohorts are provided in full in **Table S3**. **(B)** Reason for euthanasia as a proportion of cohort. ‘Age’ indicates no phenotype – mice had reached two years old. ‘Other’ indicates the development of tumours elsewhere than the mammary gland (the head/neck). Numbers of mice in each cohort are indicated. **(C)** Kaplan-Meier curves of latency of mammary tumours only. Full statistics are provided in **Table S3**. Due to the small number of mammary tumours in the *K14Cre-NeuKI* cohorts, data for the virgin and parous animals was combined. **(D)** Mean (±SD) numbers of tumours per animal in each cohort. The total number of animals assessed is indicated. Full statistical information is provided in **Table S3**. **(E)** Body location of mammary epithelial tumours in each cohort. The number of tumours assessed is indicated (which may include more than one tumour from one animal). Anterior indicates tumours developing in the 1^st^/2^nd^/3^rd^ mammary fat pads; posterior indicates tumours developing in the 4^th^/5^th^ fat pads. Right/left indicates tumours developing on the right/left hand side of the body, when viewing the animal from the dorsal aspect. **(F)** Distribution of mammary epithelial tumour histopathological phenotypes within tumour cohorts. The number of tumours in each group which underwent detailed histological analysis is indicated. AC(NST), Adenocarcinoma (No Special Type); AME, adenomyoepithelioma; ASQC, adenosquamous carcinoma. Sarcomas arising in the mammary gland and elsewhere are not included. Data for the virgin and parous *K14Cre-NeuKI* tumours is combined. Full details of all tumours is provided in **Table S4**.

Control cohort animals survived >600 days whereas both parous and virgin *MMTV-NeuNDL* mice rapidly succumbed to mammary tumours (median latencies 141 and 210 days respectively). *BlgCre-NeuKI* mice survived longer than *MMTV-Neu* mice but had a significantly shorter survival than *K14Cre-NeuKI* mice (**Figure 1A**, **Table S3**). Indeed, the majority of the latter were culled due to age rather than pathology, whereas *BlgCre-NeuKI* mice developed tumours in a number of sites, mainly the mammary gland but also on the head and neck (some *BlgCre-NeuKI* mice were also culled due to age; **Figure 1B**). This is in strong contrast to our previous studies on mice carrying conditional *Brca1*, *p53* or *Pten* alleles, in which mice carrying the *K14Cre* transgene had a significantly shorter survival than *BlgCre* mice [7, 8].

Parity significantly accelerated tumour onset in *MMTV-Neu* mice. It also lowered median overall survival and tumour-specific survival in *BlgCre* mice (median overall survival virgin animals 470 days, parous animals 245 days; tumour-specific survival virgin animals 457 days, parous animals 245 days); however, the overall survival effects were not significant and the significance of the difference in age of mammary tumour onset depended on the statistical test (**Figure 1C, Table S3**). The median tumour-specific survival of the combined *K14Cre-NeuKI* cohort was 566 days.

Therefore, whereas our previous findings established that basal mammary cells were more sensitive than luminal cells to loss of *Brca1/2*, *Pten* and *p53*, here we demonstrate that luminal cells are more sensitive to activation of the *NeuKI* allele and, indeed, that the basal cell population is largely refractory to the tumour-promoting activity of this allele.

### Parity alters tumour phenotype in the BlgCre-NeuKI model

We next focused on the numbers and locations of mammary tumours in the tumour cohorts on a mouse-by-mouse basis, as well as on the tumour phenotypes. Given the small number of mammary tumours from the *Krt14Cre-NeuKI* model, data from the virgin and parous cohorts of this line were combined for this analysis.

Mice from the *MMTV-NeuNDL* cohorts developed on average 2.2 tumours per animal, whereas *Krt14Cre-NeuKI* mice and *BlgCre-NeuKI* virgin animals developed 1.3 and 1.2 tumours per animal respectively (**Figure 1D** and **Table S3** for full statistical comparisons). There was no significant difference in mean tumour number per animal between *BlgCre-NeuKI* parous mice (1.7) and either the virgin mice of the same cohort or the *MMTV-NeuNDL* mice. However, there was a wide range in the numbers of tumours per animal seen in the *BlgCre-NeuKI* parous cohort (**Table S1**), suggests that whilst pregnancy was having a biological effect on this cohort, it was not consistent between animals.

Breast cancer shows a laterality bias, with an approximately 10% higher incidence in the left breast [27–29]. We therefore analysed mouse necropsy data for evidence of a locational bias in tumour origins. For this analysis, virgin and parous cohorts were combined and we categorised tumours as originating on either the left or right side of the mouse or in the anterior or posterior fat pads (mammary fat pads 1, 2 and 3 being anterior; 4 and 5 being posterior) (**Figure 1E**). There was no significant left-right bias in tumour origins in any of the mouse lines. There was no significant anterior-posterior bias in the *Krt14Cre-NeuKI* and *BlgCre-NeuKI* lines. However, in the *MMTV-NeuNDL* line, tumours were significantly (P<0.0001) more likely to develop from the anterior as opposed to the posterior fat pads.

For the analysis of tumour phenotypes, we undertook a detailed comparative study as previously described [7, 8]. In brief, mouse mammary tumours fall into four main histotypes: adenomyoepithelioma (AME), metaplastic adenosquamous carcinoma (ASQC), metaplastic spindle cell carcinoma (MSCC) and adenocarcinoma of no special type (AC(NST)) which can be readily diagnosed from H&E and ΔNp63 staining (which identifies key differential diagnostic features, namely the presence of metaplastic features and the number/pattern of ΔNp63 stained cells). Full details of the analysis are given in **TableS4**. Ten virgin and ten parous *MMTV-Neu* tumours were analysed. These were invariably diagnosed as AC-NST (**Figure 1F**). They grew as sheets of epithelioid cells; only one showed evidence of metaplasia. In these tumours, ERBB2 showed, as expected, extremely strong membrane staining (**Figure 2A**).

**Figure 2:**
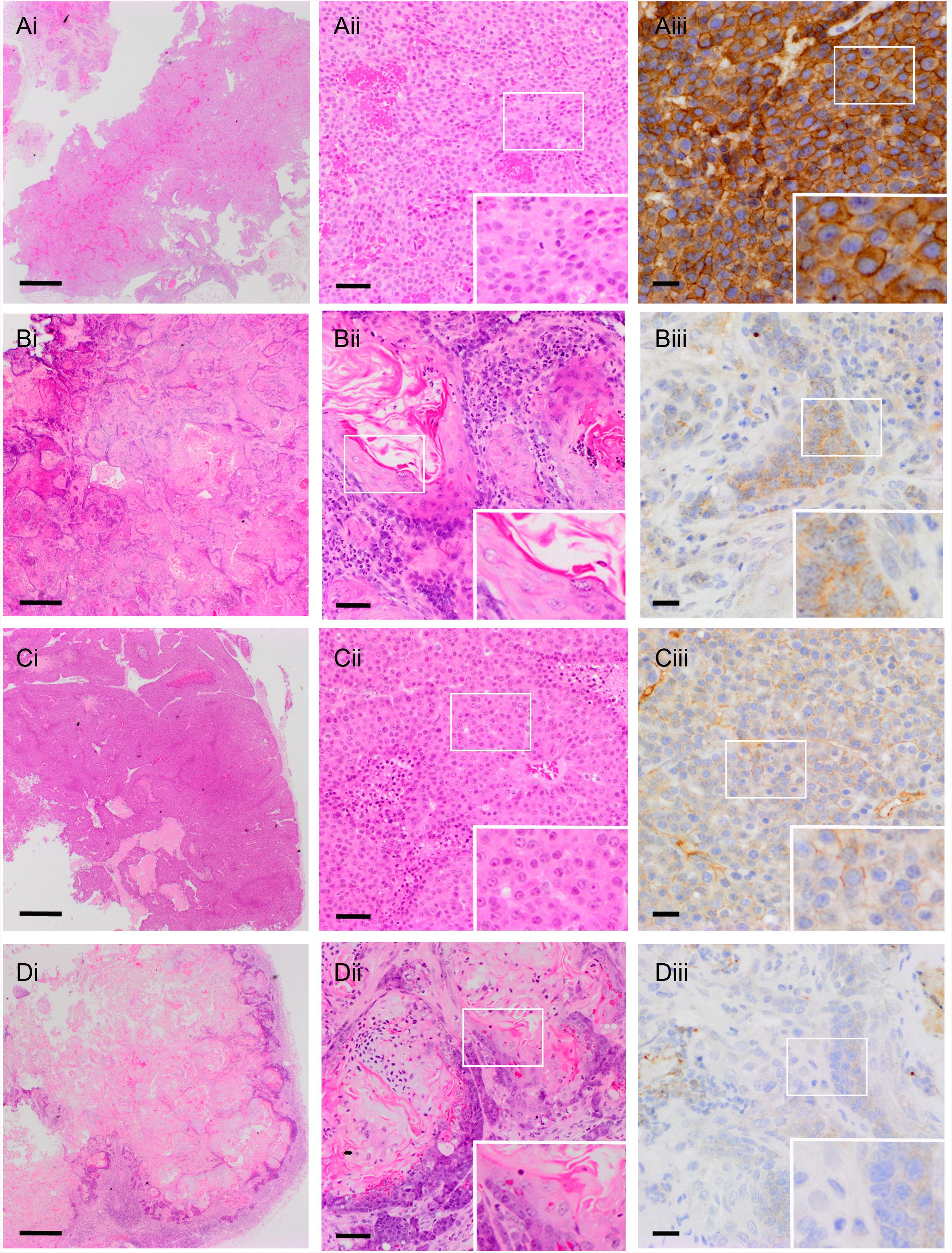
Tumour histology. Representative low (column (i); bars = 500 μm) and high power (column (ii); bars = 50 μm) H&E images and ERBB2 staining (column (iii); bars = 20 μm) of **(A)** a virgin *MMTV-NeuNDL* tumour, **(B)** a virgin *Krt14Cre-NeuKI* tumour, **(C)** a virgin *BlgCre-NeuKI* tumour and **(D)** a parous *BlgCre-NeuKI* tumour. Insets are a 2x magnification of the boxed region in each panel.

Six virgin and one parous *K14Cre-NeuKI* tumours were available for histological analysis. The low numbers resulted from the cystic nature of many of these tumours, which collapsed upon excision and left little or no material for embedding. Of those that could be analysed, five were diagnosed as AMEs and two as ASQC tumours (**Figure 1F**), consistent with our previous findings on phenotypes of tumours arising from the basal populations. ERBB2 was not membrane-localised in these tumours and in at least one tumour, staining was punctate within the cytoplasm (**Figure 2B**).

Analysis of the *BlgCre-NeuKI* tumours was the most striking. Seventeen virgin and twenty-three parous tumours arising in the mammary gland were available for analysis. One of the virgin and two of the parous tumours were diagnosed as malignant mammary sarcomas and excluded from the analysis (note, the head/neck tumours were also diagnosed as sarcomas). Of the remainder, 50% (n=8) virgin tumours were classed as AC(NST), 25% (n=4) as AMEs and 25% (n=4) as ASQCs. In contrast, 38% (n=8) parous tumours were AC(NST), 5% (n=1) was an AME, and 57% (n=12) were ASQCs. This was a statistically significant increase in ASQC tumours in parous compared to virgin *BlgCre-NeuKI* mice (**Figure 1F**). In contrast to the strong membrane staining seen in *MMTV-Neu* tumours, ERBB2 staining in *BlgCre-NeuKI* tumours was weak and typically cytoplasmic, with weak membrane staining in less than half of all cases (**Figure 2C,D**). In some tumours, ERBB2 staining was undetectable. No MSCCs were observed in any cohort.

Importantly, the parous and virgin *BlgCre-NeuKI* cohorts were established from litter mates randomly assigned to the groups and therefore of identical background genetics. Thus, in this model, developmental history of the mammary gland has a biological effect on tumour phenotype, with parity biasing towards the formation of ASQC tumours.

### *AC(NST) tumours show the largest copy number gain at the* Errb2/Neu *locus*

It was previously demonstrated that tumour formation in a mouse model in which the *<u>NeuKI</u>* allele was activated by an *MMTV-Cre* transgene was associated with amplification of this locus [9]. To test whether the *Neu* allele was also amplified in tumours from the *BlgCre NeuKI* model, we tested a subset of tumours with a quantitative PCR approach using a *Neu*-specific probe on genomic tumour DNA. The tumours consisted of a mixture of AC(NSTs) and ASQCs (9 vs 5) which had come from both virgin and parous animals (5 vs 9). In addition, one non-epithelial tumour (a mammary sarcoma) was included (**Figure 3A**). This analysis suggested that the *Erbb2/Neu* locus was typically most highly amplified in AC(NSTs) relative to control DNA from normal primary mammary cells. In contrast, in ASQC tumours and the sarcoma the locus showed lower levels of amplification, although some AC(NST)s (e.g. MS1166.1) also fell within the low amplification group.

**Figure 3:**
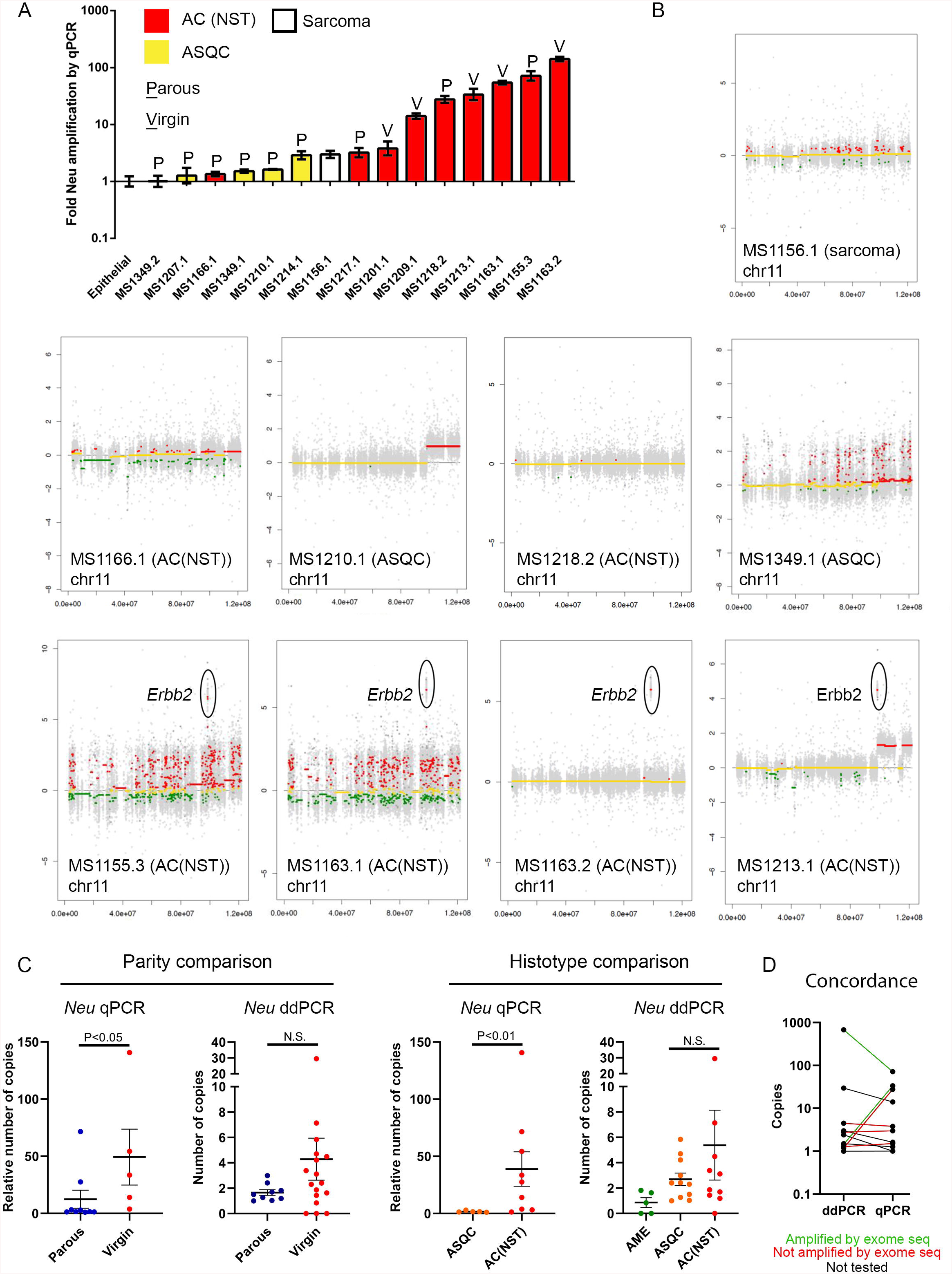
*Neu* allele amplification in *BlgCre-NeuKI* tumours. **(A)** *Neu* allele amplification assessed by qPCR on tumour genomic DNA relative to control normal primary mammary epithelial cells. Mean±95% confidence intervals of three technical replicates on each tumour. Parity status and tumour histotype are indicated. **(B)** CNV-by-exome analysis of chromosome 11 in nine tumours from *BlgCre-NeuKI* cohort. The tumour type is indicated and amplification of the *Erbb2* locus in four tumours indicated. **(C)** Comparison of Neu amplification status (mean±SEM) by parity and histotype as assessed by qPCR and ddPCR. Parity status/histotypes are indicated. P values (two-tailed t-test with Welch’s correction on Log10 transformed values). **(D)** Concordance in findings between tumours analysed for *Neu* copy number by both ddPCR and qPCR, with CNV-by-exome findings indicated by the colour of the connecting line where available. The majority of tumours have similar copy number levels by both methods, although there is more variability in tumours with the very highest values. However, two tumours score ‘low’ Neu by ddPCR but ‘high’ Neu by qPCR. One of these was found to be amplified, while the other was not amplified, by CNV-by-exome. This argues against a systematic error in the methodologies and in favour of sampling errors relating to tumour heterogeneity.

We selected ten tumours which had been copy number profiled by qPCR, together with matched spleens, for exome sequencing. The tumour numbers and their features are given in **Table S5**, detailed exome sequencing results are provided in **Table S6** and a summary of the results in **Table S7**. DNA/RNA isolated from one ASQC sample (MS1207.1) was of insufficient quality for sequencing. The remaining nine tumours consisted of six AC(NST), two ASQC tumours and a sarcoma. The nine tumours had a median of 278 mutations, both coding and non-coding (range 167 - 636). There were a median of 19 coding mutations (range 13 – 33) predicted to alter protein expression (**Table S7**). However, only 14 genes were mutated in more than one sample, and only two of these in more than two samples. Furthermore, there was no consistent association with parity or tumour phenotype (**Table S7**) and some recurrently mutated genes were identified in the sarcoma as well as epithelial tumours. These findings suggested that coding sequence mutations did not underlie the phenotypic changes seen in parous compared to virgin tumours.

Using the exome data to estimate copy number variations (CNVs) confirmed the qPCR analysis of the *Erbb2/neu* locus in genomic DNA in eight of the nine tumours (**Figure 3B**, **Table S8** and **S9**). Four tumours which the qPCR reported had a strongly amplified *Neu* oncogene, all of which were AC(NST), showed amplification by exome data of a segment in chromosome 11 which included *Erbb2* but also nearby genes *Mien1* and *Grb7*. Two tumours also had amplified loci on chromosome 19 which included *Pten*, a key negative regulator of PI3K-AKT signalling, and *Atad1*. In four tumours, two of which were ASQCs, one an AC(NST) and the mammary sarcoma, all of which had low levels of amplification by qPCR, no CNVs could be detected which passed the statistical threshold (**Figure 3B**, **Table S8** and **S9**). Only one tumour, an AC(NST) (MS1281.2) showed discordant results between the qPCR and sequencing-based analysis of CNVs.

To broaden these findings on copy number variations across a larger tumour sample, we carried out ddPCR for the genomic *Erbb2/Neu*, *Grb7*, *Mien1*, *Pten* and *Atad1* loci on the tumour cohorts and matched spleen samples, using both snap frozen tumour samples and DNA isolated from FFPE blocks. We also analysed the *NeoR loxP-stop-loxP* cassette to confirm its deletion (and thus activation of the *NeuKI* locus) in tumour samples, especially those ASQCs in which locus amplification had not occurred. We included *Erbb2/Neu* locus-specific probes to both the mutant *NeuKI* allele and exon 12 of the endogenous *Erbb2* gene (see schematic of the locus in **Figure S1**). As controls, DNA extracted from primary mammary epithelial organoids isolated from 10-week-old virgin wild type mice, or mice carrying the *NeuKI* allele but no CRE, and cultured *in vitro* for 7 days were used.

As expected, in wild-type organoids *NeoR* and *Neu* were undetectable whereas *Erbb2* exon 12 was present in two copies. In *NeuKI* organoids, as expected, *NeoR* and *Neu* were each present in a single copy and *Erbb2* exon 12 was present in two copies. In the majority of spleens from tumour cohort animals (**Figure S2A**), *NeoR* and *Neu* were also present in a single copy, and *Erbb2* in two copies, again as expected as *Blg* is considered a mammary-specific promoter. However, in spleens from two animals, MS1163 and MS1379, the *NeoR* cassette had been lost and *Neu* was amplified. In MS1379, *Erbb2* was also amplified. Therefore, there is a low level of background recombination in this allele in the spleen, either due to leakiness of the *Blg* promoter or non-specific effects of endogenous recombinases in the spleen.

26 out of 29 tumours tested (of all phenotypes) showed evidence of *NeoR* recombination, or locus amplification, or both (**Figures S2B-E)**. The exceptions were three ASQC tumours. When comparing tumours either by the parity status of the animals or the tumour histotype, there was broad agreement in the findings from the ddPCR with the CNV-by-exome analysis and the original genomic qPCR analysis of the *NeuNT/Erbb2* locus (**Figure 3C; Table S10).** The mean copy number of the *Neu* allele was higher in tumours from virgin animals, and in tumours of the AC(NST) histotype. However, the range of amplification was wide and these differences were significant by genomic qPCR but not significant by ddPCR for both comparisons. Furthermore, when considering individual tumours in which both ddPCR and qPCR had been performed on the *Neu* allele, there were some discordant results (**Figure 3D**). It is not clear whether the differences arising from different approaches, and the discordant results between some individual tumours, are a result of technical limitations or tumour heterogeneity leading to sampling biases. However, all three approaches used to analyse *NeuKI* allele copy number support the model that AC(NST) tumours tend to have higher levels of *NeuKI* amplification, whilst ASQC tumours have lower levels.

Confirming the CNV-by-exome analysis, ddPCR analysis of *Grb7* and *Mien1*, located close to *Erbb2* in both mouse and human (the locus is syntenic in the two species), demonstrated similar levels of amplification in the tumours to *Erbb2/Neu* (**Figure S3; Table S10**). This supports a model that the whole locus is a hotspot for amplification, and it is not simply a feature of the engineered allele. In contrast, while amplification of *Atad1* and *Pten*, as suggested by the CNV-by-exome analysis, was confirmed by ddPCR in some tumours, potential copy number losses were also observed in others (**Figure S4; Table S10**). Thus, the significance of changes in *Pten* and *Atad1* is uncertain.

### Erbb2/Neu *locus amplification is the prime determinant of tumour phenotype and distinct gene expression patterns are associated with locus amplification state*

Next, we carried out RNAseq analysis on the same tumours used for exome sequencing. Data was compared across the tumour sets as a whole to identify genes that were differentially expressed between biologically distinct tumour groups. Non-negative matrix factorisation (**Methods**) was used to compare samples on the basis of parity, tumour phenotype and *Erbb2/Neu* locus amplification status (by genomic qPCR) and to determine which of these comparisons generated the most distinct and stable sample clusters i.e. which was the strongest driver of differences in gene expression pattern. The results showed that the strongest determinant of gene expression differences between the samples was amplification status of the *Erbb2/Neu* locus in the epithelial tumours, with the mammary sarcoma being an outlier (**Figure 4A** and **Figure S5**).

**Figure 4:**
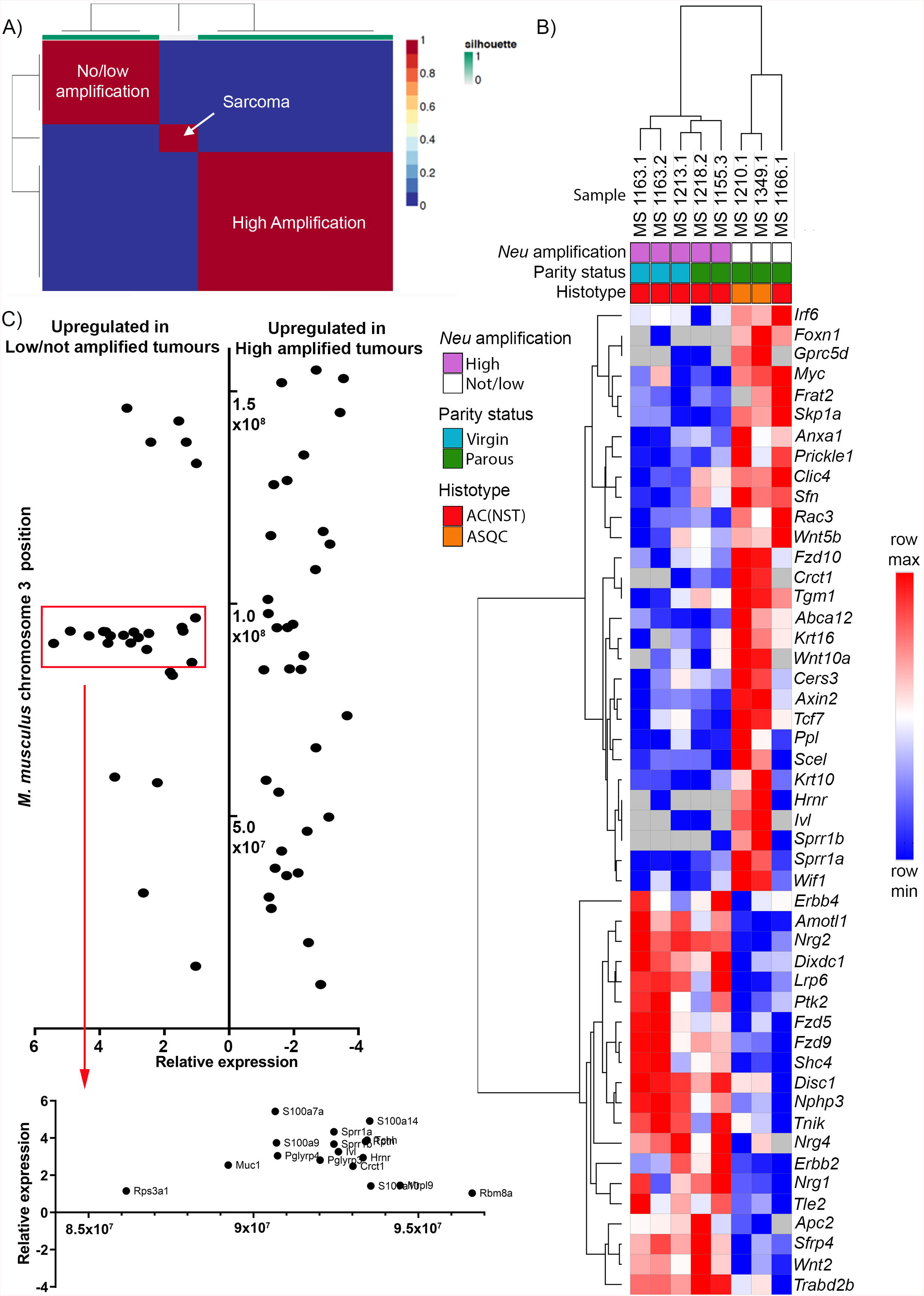
*Neu* amplification status is associated with canonical and non-canonical WNT signalling and activation of the Epidermal Differentiation Cluster. **(A)** Cluster allocation analysis of RNASeq data using Non-negative Matrix Factorisation to identify correlations between tumour characteristics and gene expression. **(B)** Unsupervised hierarchical clustering of Log2 RPKM data for genes of interest identified by functional annotation (from GO/KEGG terms GO:0016055~Wnt signalling pathway, GO:0060070~canonical Wnt signalling pathway, GO:0090263~positive regulation of canonical Wnt signalling pathway, GO:0030216~keratinocyte differentiation, GO:0031424~keratinization, mmu04310:Wnt signalling pathway and mmu04012:ErbB signalling pathway) (see **Table S13**). Tumours cluster by activation of canonical/non-canonical WNT signalling pathways. **(C)** Log2 relative expression of genes on *M. musculus* chromosome 3 significantly differentially expressed between low/not amplified and high amplified tumours. Genes are plotted by their relative expression levels (x-axis) and their chromosomal location (y-axis) (**Table S14**). Note the cluster of genes highly expressed in the low/not amplified tumours just before base position 1×10^8^ (red rectangle) suggesting co-regulation. This region is expanded below and the genes annotated. This is the Epidermal Differentiation Cluster, and includes genes associated with epithelial keratinisation.

As amplification status was the strongest determinant of gene expression differences, the RNAseq data was analysed to determine which genes were significantly differentially expressed between amplified (n=5; all AC(NST)) and non-amplified (n=3; two ASQC and one AC(NST)) tumours (excluding the data from the sarcoma). Normalised RPKM values for each of these tumours are shown in **Table S11** and the full list of genes significantly differentially expressed between the two classes is given in **Table S12**. 865 genes were significantly expressed >2 fold higher in amplified vs non-amplified tumours; with 619 genes being significantly >2-fold higher in the non-amplified tumours vs amplified tumours.

Functional annotation and functional annotation clustering of differentially expressed genes was carried out using the DAVID (v6.8) on-line tool [30, 31] on the basis of bioprocess Gene Ontology and KEGG pathway analysis (**Table S13**). Functional annotation clustering of the amplified tumour genes indicated that these tumours were enriched for protein glycosylation, protein phosphorylation, genes associated with auditory morphogenesis and multiple signalling pathways including most notably ERBB signalling, PI3K-AKT signalling and WNT signalling. The not/low amplified tumours were enriched for fatty acid biosynthetic process, cell division/cell cycle, regulation of transcription, cytokine signalling/chemotaxis, keratinisation and skin development and also, interestingly, WNT signalling.

The enrichment of amplified tumours for ERBB signalling was consistent with the amplification of the *Neu* locus and the enrichment of non/low-amplified tumours for keratinisation and skin development was consistent with their enrichment for ASQC phenotype tumours (even allowing for the fact that of the three non/low-amplified tumours available for RNAseq analysis, only two were of ASQC phenotype). However, the finding that both groups of tumours were enriched for WNT signalling was intriguing. Notably, the two tumour classes were enriched for a different set of WNT-associated genes (amplified tumours, *Fzd9*, *Dixdc1*, *Tnik*, *Apc2*, *Trabd2b*, *Tle2*, *Amotl1*, *Fzd5*, *Wnt2*, *Nphp3*, *Sfrp4*, *Lrp6*, *Disc1*; non/low-amplified tumours, *Wnt10a*, *Fzd10*, *Tcf7*, *Wnt5b*, *Prickle1*, *Rac3*, *Frat2*, *Wif1*, *Skp1a*, *Axin2*, *Myc*). To better understand how the WNT, ERBB and keratinisation genes defined the differences between the two tumour classes, the individual RPKM values of the set of 49 genes from the GO bioprocess/KEGG pathway terms GO:0016055~Wnt signalling pathway, GO:0060070~canonical Wnt signalling pathway, GO:0090263~positive regulation of canonical Wnt signalling pathway, GO:0030216~keratinocyte differentiation, GO:0031424~keratinization, mmu04310:Wnt signalling pathway and mmu04012:ErbB signalling pathway were analysed by unsupervised hierarchical clustering (**Figure 4B**). As expected, clustering using these genes divided the tumours into two main groups based on amplification status. Furthermore, the two different sets of WNT-associated genes clearly differentiate between the two tumour classes.

### The Epidermal Differentiation Cluster is activated in ASQC tumours

The non/low-amplified tumours showed upregulation of keratinisation-associated genes although only two of them displayed a clear ASQC phenotype. Keratinisation is driven by activation of the Epidermal Differentiation Cluster (EDC) [32], a group of co-regulated epidermal genes. To determine whether there was evidence for activation of the EDC in non/low-amplified tumours/ASQCs, the genomic locations of each of the 1,484 significantly differentially expressed genes from **Table S12** were retrieved from the JAX Mouse Genome Informatics database, taking the chromosome on which the gene is located and its ‘start position’ as the genomic location (**Table S14**).

In mice, the EDC is located on chromosome 3. Therefore, the relative expression levels of the differentially expressed genes located on chromosome 3 were plotted against their genomic locations (**Figure 4C**). Notably, the genes on this chromosome which are upregulated in the low/not amplified tumours cluster to the EDC in a location just proximal to the 1 × 10^9^ base position, supporting the model that squamous metaplasia in ASQC tumours is driven by co-ordinated activation of genes of the EDC.

### ASQC tumours have activated canonical WNT signalling

The WNT-pathway associated genes differentially expressed between not/low amplified and high-amplified tumours suggest different branches of the WNT signalling pathway were being activated in these tumours. The two main WNT pathways are the canonical signalling pathway, characterised by the role of β-catenin as a nuclear transcription factor driving gene expression changes, and the non-canonical or planar cell polarity pathway, which regulates the actin cytoskeleton and is an important regulator of collective cell migration in tumour metastasis [33]. The WNT ligands expressed at high levels in the non/low-amplified tumours (WNT5B, WNT10A) are associated with canonical signalling, while WNT2 (expressed in the high amplified tumours) is associated with non-canonical signalling.

To confirm whether canonical WNT signalling was activated specifically in ASQC tumours, β-catenin localisation was assessed in AC(NST) and ASQC tumours from the *BlgCre-NeuKI* tumour cohort (**Figure 5A, B**). Nuclear β-catenin staining was highly significantly (P<0.001) associated with ASQC tumours, although there was no significant difference in levels of *Ctnnb1* gene transcription between the histotypes (**Figure 5C**). Next, expression levels of four well-characterised canonical WNT target genes (*c-Myc*, *Tcf7*, *Axin2*, *Wif1*) were compared between the two histotypes by real time qrtPCR (**Figure 5D**). There was no difference in *c-Myc* expression between AC(NST) and ASQC tumours, however, expression of *Tcf7*, *Axin2* and *Wif1* were all significantly (p<0.01) elevated in ASQC tumours. Canonical WNT signalling is, therefore, activated in the ASQC histotype.

**Figure 5:**
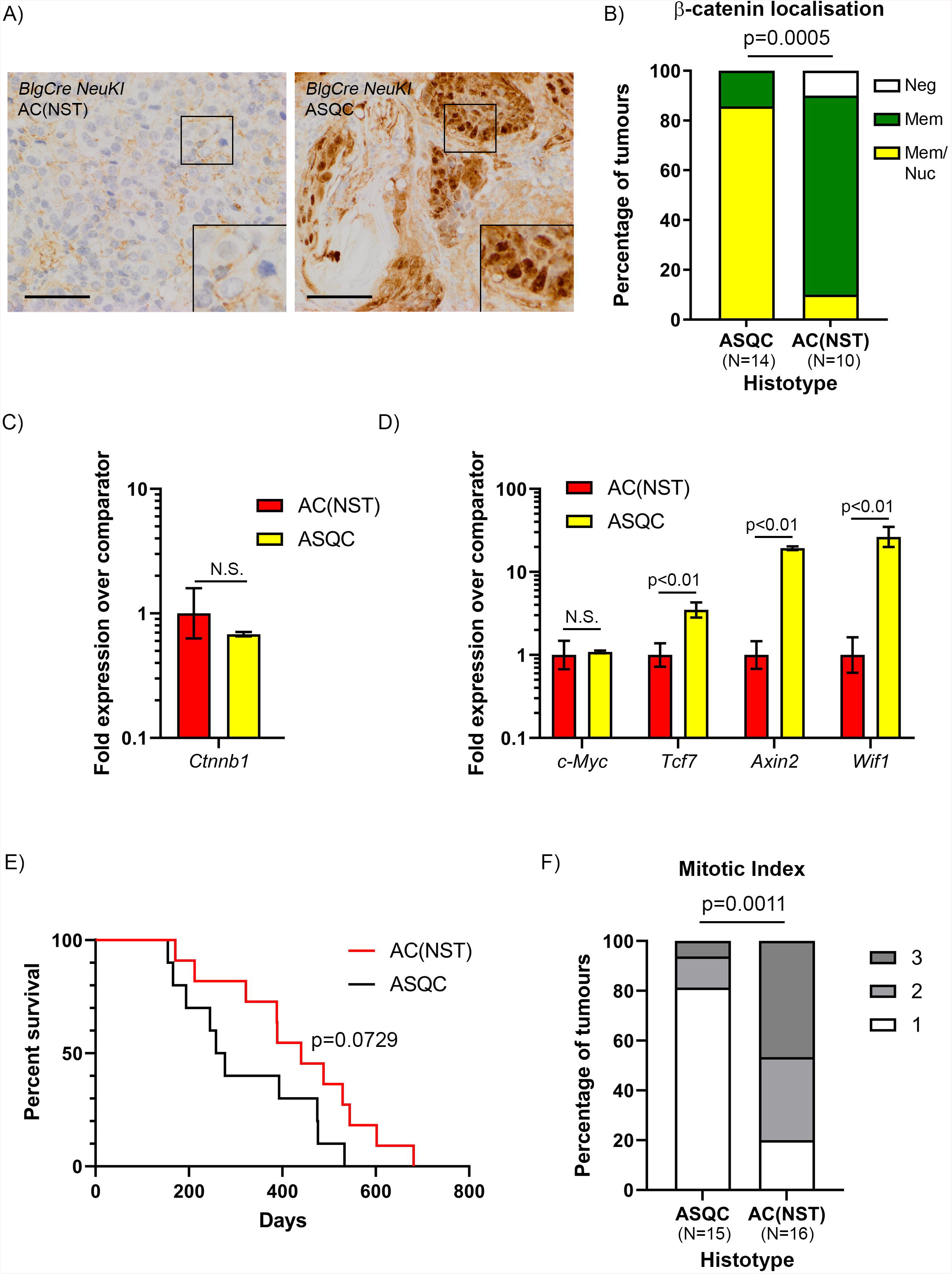
Canonical WNT signalling is activated in ASQC tumours. **(A)** Representative examples of membrane and nuclear β-catenin localisation in AC(NST) and ASQC tumours. Bars = 50 μm. **(B)** Quantitation of β-catenin localisation. p=0.0005, Fisher’s exact test of proportion of tumours with nuclear staining vs no nuclear staining/no staining). **(C)** Real time qrtPCR analysis of *Ctnnb1* gene expression. Data presented as mean fold expression±95% confidence intervals relative to comparator (the AC(NST) cohort) (N = eight AC(NST) and six ASQC tumours, three technical replicates of each tumour; significance determined from confidence intervals according to [26]). **(D)** Real time qrtPCR analysis of canonical WNT target gene (*c-Myc, Tcf7, Axin2, Wif1*) expression. Data presented as mean fold expression±95% confidence intervals relative to comparator (the AC(NST) cohort) (N = eight AC(NST) and six ASQC tumours, three technical replicates of each tumour; significance determined from confidence intervals according to [26]). **(E)** Kaplan-Meier survival curves for *BlgCre NeuKI* mice according to tumour histotype. Only mice who has one or more ASQC tumours (N = 10 animals), or one or more AC(NST) tumours (N = 11 animals) were included. Mice with AME tumours, non-mammary epithelial tumours or who developed multiple tumours of different histotypes were not included. P=0.0729 by Log-Rank test. (F) Comparison of mitotic index in 15 ASQC and 16 AC(NST) tumours from *BlgCre NeuKI* mice. Fisher’s exact test of proportion of tumours with grade 1 mitotic index vs tumours with grade 2 or 3 mitotic index as defined the WHO breast cancer classification criteria [11].

To assess the significance of elevated expression of canonical WNT target genes in human HER2 breast cancer, the KM plotter resource [34] was used to mine relapse-free survival data for unselected human breast cancers, for HER2-amplified breast cancers and for HER2-non-amplified breast cancers stratified according to *c-MYC*, *TCF7*, *AXIN2* and *WIF1* expression (**Figure S6**). The results show that high expression of *TCF7*, *AXIN2* and *WIF1* all predict shorter Relapse-Free Survival (RFS) in HER2-amplified breast cancer, but longer RFS in HER2-non-amplified disease. Interestingly, the pattern is reversed for *c-MYC* expression. Although RFS in human disease cannot be directly correlated with survival curves for mouse tumour models, these findings suggested that expression of canonical WNT target genes in the mouse tumours might be associated with differences in survival. We had already shown that parity in the *BlgCre-NeuKI* cohort was associated with a slightly shorter median survival compared to virgin animals, although this was of borderline significance (**Figure 1C**). We had also shown the ASQC phenotype (in which canonical WNT signalling is active) was associated with parity (**Figure 1F**). Therefore, we directly tested whether animals which developed only ASQC phenotype tumours had a difference in survival compared with animals which developed only AC(NST) tumours. Survival curves based solely on tumour phenotype showed that ASQC tumour-bearing animals had a shorter median survival of 267 days compared to AC(NST) animals with a median survival of 440 days (**Figure 5E**) although this was not significant. However, it is important to note that these curves are based on the time at which mice had to be euthanised because the tumours reached specified size limits. They do not reflect growth rates of the tumours. Indeed, comparison of mitotic index scores, based on the WHO breast tumour grading guidelines, demonstrate that ASQC tumours had a significantly lower mitotic index than AC(NST) tumours (**Figure 5F**). This suggests that ASQC tumours had an earlier onset than AC(NST) tumours but grew more slowly, therefore the overall survival of the animals with the tumours was not significantly different.

Therefore, unlike our previous analyses of the sensitivity of different mammary epithelial cell populations to loss of *Brca1*, *Brca2*, *Pten* and *p53*, the basal mammary epithelial layer is resistant to transformation by activation of the *NeuKI* allele, whereas the luminal ER negative mammary epithelial cells are sensitive. Activation of the allele in luminal cells in virgin animals tends to generate AC(NST) tumours associated with high locus amplification and activation of non-canonical WNT signalling. In contrast, in parous animals, activation tends to generate ASQC tumours with lower levels of locus amplification and activation of both the Epidermal Differentiation Cluster and canonical WNT signalling. ASQC tumours tend to occur earlier than AC(NST) tumours but grow more slowly. The key features of the different models we describe here are summarised in **Table 1**.

**Table 1:** Comparison of main features of *NeuKI* tumour models. AC(NST), Adenocarcinoma (No Special Type); AME, Adenomyoepithelioma; ASQC, Adenosquamous Carcinoma.

| Comparison | *K14Cre NeuKI* | *BlgCre NeuKI* virgin | *BlgCre NeuKI* parous | Comment |
| --- | --- | --- | --- | --- |
| Median overall survival (days) | Could not be defined | 470 | 245 | **Figure 1A**; see **Table S3** for statistical comparisons. |
| Tumour penetrance (all sites; % of animals) | 37 | 83 | 84 | **Figure 1B**; see **Table S3** for statistical comparisons. |
| Median tumour-specific survival (days) | 566 | 457 | 245 | **Figure 1C**; see **Table S3** for statistical comparisons. |
| Tumour phenotypes (%) | AME: 71<br>ASQC: 29<br>AC(NST): 0 | AME: 25<br>ASQC: 25<br>AC(NST): 50 | AME: 5<br>ASQC: 57<br>AC(NST): 38 | **Figure 1F**; human HER2 breast cancers are typically equivalent to AC(NST) [11] |
| Mean *NeuKI* allele copy number (qPCR) | N/D | 49.29 | 12.45 | **Figure 3C** |
| Mean *NeuKI* allele copy number (ddPCR) | N/D | 4.29 | 1.66 | **Figure 3C** |
| Comparison |  | *NeuKI* High amplification | *NeuKI* Low amplification | Comment |
| GO/KEGG enriched processes/pathways |  | protein glycosylation, protein phosphorylation, auditory morphogenesis, ERBB signalling, PI3K-AKT signalling, WNT signalling | fatty acid biosynthetic process, cell division/cell cycle, regulation of transcription, cytokine signalling/chemotaxis, keratinisation and skin development, WNT signalling | **Table S13** |
| Activation of Epidermal Differentiation Cluster |  | No | Yes | **Figure 4C** |
| WNT signalling |  | Non-canonical | Canonical | **Figure 5** |

## Discussion

Use of *MMTV*-*Neu* mouse models has been key in identifying many of the genetic and molecular events that control *HER2*-induced breast cancer progression [35, 36]. Both wild type (*NeuN*) and activated (*NeuNT*) forms of Neu have been used, however, both drive rapid tumour formation as a result of the activity of a strong exogenous promoter, rather than amplification of the genomic locus, as occurs in the human disease [37]. In contrast, when *NeuNT* was knocked into the endogenous *Erbb2* locus, and therefore expressed under the control of the endogenous promoter, *MMTV-Cre*-dependent activation of expression resulted in mammary hyperplasia and the formation of focal tumours after a long latency (over 1 year) [9]. Importantly, tumour formation in this system was associated with amplification of the *Erbb2/NeuNT* locus [9] by an unknown mechanism. Tumours from the *MMTV-Cre NeuNT knock-in (MMTV-Cre NeuKI)* mouse model showed similar additional genomic abnormalities to *HER2*-initiated human breast cancer, including centrosome abnormalities and recurrent deletions of chromosome 4 [38, 39]. However, the target cell for tumour formation in both *MMTV-NeuN/NT* and *MMTV-Cre NeuKI* models has not been defined and likely depends on which mouse line is used. These models have been unable, therefore, to contribute to our understanding of how cell-of-origin may affect HER2 amplification, tumour formation, phenotype and behaviour in humans.

We have previously demonstrated that tumour phenotypic heterogeneity is driven by a combination of the cell-of-origin and initiating genetic lesion, and that the ER− luminal stem/progenitor population has the potential to be the origin of both TNBC-like and ER positive-like mammary tumours. To determine whether cell-of-origin also affects development and phenotype of HER2-amplified tumours, we used our cell-type-targeted promoter approach to activate the *NeuNT* allele in either basal or luminal ER− mammary epithelial populations. In contrast to our previous studies in which *Brca1*, *Brca2*, *Pten* and *p53* were conditionally deleted in these populations [7, 8], we found that the basal mammary epithelium is refractory to transformation by *NeuKI* activation. This demonstrates that at least one reason for distinct organ tumour tropisms associated with specific somatic or germline mutations is that individual cell types are differentially sensitive to transformation by different genetic lesions. The reason that luminal ER− cells are sensitive to *NeuKI* activation is not clear, but we suggest it is at least in part due to the role of WNT signalling in mammary development/pregnancy and interactions between these two pathways.

The complex relationship between WNT signalling and mammary development is a result of the activation of either canonical or non-canonical WNT signalling at different stages of the developmental cycle of the mammary gland. During early mammary development and the ductal elongation which occurs during puberty, the most highly expressed mammary WNTs are WNT2 and WNT5A (which are associated with activation of non-canonical WNT signalling) [40]. This is consistent with the role of non-canonical WNTs in planar cell polarity and the regulation of collective cell migration [33]. However, during pregnancy there is a switch to the activation of canonical WNTs, in particular WNT4, WNT5B and WNT6 [40]. The canonical pathway is typically associated with regulation of stem/progenitor cells, consistent with the activation of alveolar progenitors and the remodelling of the gland for milk production. Notably, we have previously demonstrated that strong activation of WNT signalling in organoid culture *in vitro* (by R-spondins) results in squamous metaplasia [41]. Here, our *in vivo* findings support this, with the ASQC tumour phenotype strongly associated with activation of canonical WNT signalling but with the trigger in this case being pregnancy. Pregnancy results in a spike of canonical WNT signalling within the mammary gland. Our findings suggest that in the context of activation (but not amplification) of the *NeuKI* allele, this canonical WNT spike persists and becomes a chronic activity that leads to formation of tumours with squamous metaplasia. We suggest that in this system the canonical WNT and NeuKI allele-driven pathways are interacting at the level of β-catenin and FoxO factors.

Transcription factors of the Forkhead class (FoxO) have been characterised as positive transcriptional regulators of pro-apoptotic (such as BIM) and cell cycle arrest (such as p27) genes [42]. In response to activation of PI3K-AKT signalling, nuclear FoxOs are phosphorylated and as a result translocated out of the nucleus. Apoptosis is thus suppressed and cell cycle inhibition relieved, allowing proliferation. We propose a model in which activation of PI3K-AKT signalling by the *NeuNT* allele results in FoxO phosphorylation, translocation out of the nucleus, an increase in proliferation and protection from apoptosis. The stronger the NeuNT – PI3K – AKT signal, the more FoxO is excluded from the nucleus and the greater the increase in proliferation and protection from apoptosis. Hence, in a population of transformed cells, the greatest ‘fitness’ will be conferred on the clone with the strongest activity of the NeuNT – PI3K – AKT pathway. Therefore, clones in which the *NeuNT* locus has been amplified will emerge and dominate the tumour – the situation in virgin/amplified/AC(NST) tumours.

The interaction of WNT and FoxO signalling is not straightforward. In general, they are considered to be mutually inhibitory, antagonistic pathways. For example, canonical WNT pathway activity shifts FOXO1 from a nuclear to a cytoplasmic location in an AKT dependent manner, although how AKT is activated by canonical ligand activity remains unclear [43]. However, in colon cancer it has been reported that nuclear β-catenin and nuclear FoxO cooperate to promote tumour invasion and metastasis. Indeed, it was observed that if AKT activity was inhibited pharmacologically, the resultant increase in nuclear FoxO protein promoted cell invasion. Thus, in the context of an active canonical WNT signal, nuclear FoxO can switch from being a tumour suppressing factor to a tumour promoting factor. Therefore, the genomic variations observed in our mouse model are likely to result from changes in selection pressure on clonal populations arising from differences in the order in which WNT and NeuNT signalling are activated. If strong canonical WNT signalling occurs (for instance during pregnancy) when NeuNT has been activated but prior to expansion of a *NeuNT*-amplified clone, this would allow tumour-promoting levels of β-catenin and FoxO to accumulate in the nucleus. In this case, if *Neu* were amplified it would boost the PI3K – AKT signal, drive FoxO out of the nucleus and, in the context of canonical WNT signalling, decrease fitness. Thus, in this context, expansion of a highly amplified clone would be selected against.

Although we suggest that the emergence of *NeuNT* amplified or non/low-amplified clones depends on the outcome of interaction between WNT and FoxO signalling, it is unclear from our findings whether the WNT signal is itself directly driving locus amplification. A likely set of events which leads to *Erbb2/Neu* amplification is the acquisition of an activating mutation which leads to oncogene-induced replicative stress. This is characterised by inappropriate replication origin licencing / firing leading to, for example, collisions between the replication fork and transcriptional machinery, leading to replication fork stalling and collapse and the formation of double-stranded DNA breaks [44]. This increase in genomic instability results in junctions forming between chromosomes which resolve during cell division to create ‘neochromosomes’ – so-called chromoanasynthesis events – which are susceptible to focal amplification. Chromoanasynthesis has been demonstrated to be a mechanism of *ERBB2* amplification [45].

In the original report describing the *NeuKI* knock-in model, the *NeuKI* allele was activated by an *MMTV-Cre* promoter [9]. It is likely, therefore, that in this situation tumours had a different cell-of-origin to *BlgCre-*driven tumours. In that report, six tumours, three from virgin and three from multiparous animals, were examined for amplification of the *NeuKI* allele relative to the wild type, by Southern blot. Amplification levels were 4.8, 6 and 8.7 for parous animals and 1.7, 6.1 and 21.6 for virgin animals [9]. The histology of these tumours was not described. However, in a follow-up report, although the tumours that developed were not formally histologically subtyped they were reported as being composed of mixed cell lineages and expressing ΔNp63. The image of ΔNp63 staining presented in the report is typical of an adenomyoepithelioma [46]. This follow-up study also suggested that ERBB2 was regulated by a β-catenin/CBP complex and found that tumours had activated a WNT signalling transcriptional program and 5/20 tumours examined had nuclear β-catenin. It was not reported whether the tumours used were from virgin or parous animals. In the mouse, adenomyoepithelioma and adenosquamous tumours are typically associated with each other and with activation of the PI3K pathway [7, 47], or WNT signalling [48], or a cell-of-origin in the basal mammary epithelium [7, 8]. Therefore, the association of an AME histotype tumour with canonical WNT signalling is consistent with the findings we report here of a link between ASQC tumours and canonical WNT signalling.

A distinct feature of ASQC tumours is activation of the epidermal differentiation cluster (EDC), a group of co-regulated genes for proteins required for keratinisation of skin cells to form the protective barrier of the epidermis. Co-regulated gene clusters are typically under the control of ‘super-enhancer’ genetic elements and an H3K27ac ChIP-seq-based catalogue identified super-enhancers associated with epidermal development in the mammary epithelium [49]. In keratinocyte precursors and differentiated keratinocytes, super-enhancer elements are highly enriched in ΔNp63 binding sites [50]. ΔNp63 is a key transcriptional regulator typically expressed in basal layers of stratified epithelia, such as the epidermis, the prostatic epithelium and, in the mammary gland, the myoepithelial cells. Expression of ΔNp63 is regulated by a complex network of well-known developmental pathways (in particular NOTCH, canonical WNT, Hedgehog, FGFR2 and EGFR signalling), often with complex negative and positive feedback loops, characteristic of systems which establish and maintain tissue boundaries (reviewed in [51]). ΔNp63 expression is one of the diagnostic features of metaplastic adenosquamous tumours in the mouse and also in the human [52], with strong ΔNp63 nuclear positivity seen in regions undergoing squamous metaplasia. Thus, we suggest that in our system canonical WNT and ERBB2 pathway activation combine to activate ΔNp63, which in turn activates the EDC and drives the squamous metaplastic phenotype. It may be that differences in the regulation of ΔNp63 between the mammary epithelium of the mouse and human underly the differences in frequency of breast tumours with adenosquamous features between the two species.

Although adenosquamous tumours of the breast are rare in humans, tumours with a squamous histotype are common in other body sites, e.g. the lung, prostate, pancreas and skin. Notably, dysregulated WNT signalling has been previously linked with over-proliferative skin diseases of humans such as psoriasis [53] and activating WNT pathway mutations are important tumour drivers in human squamous cell carcinoma (SCC) [54]. Furthermore, upregulation of putative WNT transcriptional targets has been demonstrated in feline oral SCCs [55] and the murine SCC model is WNT driven [56]. Therefore, it is likely that the WNT – ΔNp63 – EDC – squamous metaplasia pathway is common across cancers and that tumours with a squamous phenotype should be considered for therapy which targets canonical WNT signalling. Notably, a recent study found that 44% of breast cancers showing squamous metaplasia had WNT pathway mutations, compared to 28% of triple negative breast cancers of no special type [57].

In human HER2-non-amplified tumours, high expression of canonical WNT target genes was associated with better prognosis. This is consistent with the less aggressive phenotype of the non-*NeuKI* amplified, canonical WNT active ASQC mouse tumours. Metaplastic adenosquamous human breast tumours, as a subset of Triple Negative Breast Cancer, would be expected to fall within the HER2-non-amplified tumour definition, although their overall rarity means they would only make up a very small proportion of that group. In contrast, canonical WNT signalling was associated with more aggressive tumours in HER2-amplified disease in humans, whereas canonical WNT signalling was not upregulated in the HER2-amplified mouse tumours. This difference highlights the *caveats* of comparing mouse and human tumours and the importance of recognising that there are likely to be both similarities and differences in the fundamental biological mechanisms driving tumour formation in the two species.

The formation of sarcomas, both in the mammary gland and on the head/neck, was a unique feature of this model. The *BlgCre* allele used here was the same as we have previously used, with no evidence of any tumour formation outside the mammary gland until now. It is notable that our analysis of *NeuKI* recombination in the spleen identified a low-level background of recombination events, and it is possible that this occurs throughout the body. Our original analysis of the mammary epithelial cell types targeted by the *BlgCre* transgene demonstrated that in an aged (42-week) virgin *BlgCre R26R* reporter mouse, 42% of luminal ER− mammary epithelial cell progenitors had undergone recombination, while 1.7% of basal progenitors and 3% of luminal ER+ progenitors had undergone recombination [8]. This biases recombination and tumour formation to the luminal ER− population but cannot exclude low-level off-target activation in the mammary epithelium and, potentially, in the wider body. If a particular off-target cell type is also particularly sensitive to the genetic lesion being induced in this manner, then a low-level background of tumours from other tissues will be seen. Potentially, mammary and head/neck mesenchymal cells are sensitive to activation of the *NeuKI* allele, but not to the other alleles we have previously used. Importantly, however, the *K14Cre* and *BlgCre* drivers we have used here are the same ones we have used in previous studies and therefore our results – in particular the difference in *K14Cre*-driven cohorts – are comparable.

In summary, targeting of *Erbb2/Neu* activation to specific cell populations within the mouse mammary epithelium supports the role of luminal ER− stem/progenitor cells as the cells of origin for most breast cancer subtypes – TNBC/’basal-like’, luminal ER+ [7, 8] and now HER2+. In this model, reproductive history, *Erbb2/Neu* activation and WNT signalling all interact to drive tumour phenotype, and a key determinant of that phenotype is the activation status of the ΔNp63-regulated Epidermal Differentiation Cluster, which underlies squamous metaplasia. These results add reproductive history to cell-of-origin and initiating genetic lesion as interacting factors which determine mammary tumour phenotypic heterogeneity. Importantly, our findings also suggest that cell-type-specific intrinsic sensitivity towards the transforming potential of activated oncogenes (and potential tumour suppressor genes) is at least one mechanism underlying the propensity for different mutations to generate tumours in a specific range of tissues.

## Acknowledgements

The authors would like to thank Prof. Barry Gusterson for support and advice on histopathological diagnosis. The authors would also like to thank the Cardiff School of Biosciences Imaging Hub for technical support in histological sample processing. We also thank Wales Gene Park (funded by Health and Care Research Wales) for infrastructure support and additional bioinformatics advice provided by Dr Kevin Ashelford. This study was supported by Cancer Research UK (C45684/A15938), Breast Cancer Now and by a Marie-Curie Fellowship to LM. LM was funded by Fundación Ramón Areces and Marie Curie Actions (IEF-236788).

This study is deposited as a preprint at bioRXiv (BIORXIV/2020/361998).

## Contributions

Conceptualisation MJS, LM; Investigation LDO, LM, KRG, HK, GT; Formal analysis LDO, LM, MJS, JB, PG, KA, KR; Writing – Original Draft, MJS, LDO; Supervision, MJS; Project Administration MJS, LDO. LM, KRG; Funding Acquisition MJS, LM.

