## Supplemental Information for "Reproductive history determines *ErbB2* locus amplification, WNT signalling and tumour phenotype in a murine breast cancer model"

**Supplementary Information**

**Supplementary Figures**

**Figure S1: Schematic of *Erbb2* locus in *NeuKI* mice.** Modified from [9] and indicating structure of wild type and engineered alleles and location of binding regions for ddPCR probes to the endogenous Erbb2 sequences (exon 12, green), the activated rat *NeuNT* sequence (blue) and the *loxP – NeoR – loxP* (STOP) cassette (red).

**Figure S2: Amplification of both *Neu* and endogenous *Erbb2* alleles in *BlgCre-NeuKI* tumours. (A – E)** Copy number of *NeoR* cassette, *Neu* and *Erbb2* alleles (mean±95% confidence intervals of three technical repeats for each sample) in spleens and tumour samples from *BlgCre-NeuKI* mice and in wild type and *NeuKI* organoid controls (eight biological replicates each of three technical replicates) by ddPCR. Samples are arranged from lowest to highest amplification from left to right. Black, *NeoR*; red, *Neu*; blue, *Erbb2*. Vertical dotted lines indicate the set of three results from each tumour sample. Horizontal lines are set at 0.5, 1.5 and 2.5. The non-recombined, non-amplified level for *NeoR* and *Neu* should be 1 (between the 0.5 and 1.5 marks); the level for *Erbb2* should be 2 (between the 1.5 and 2.5 marks). **(A)** Spleen. **(B)** Sarcomas. **(C)** AMEs. **(D)** ASQCs. **(E)** AC(NSTs). *The Neu ddPCR for MS1155-3 failed to be consistent, giving widely varying and often very high values. This result has been excluded. *Indicates analysis carried out on DNA extracted from FFPE material. For some tumours, both snap frozen and FFPE material were tested.

**Figure S3: Copy number changes in *Grb7* and *Mien1* in *BlgCre-NeuKI* tumours.** Copy number of *Grb7* (top) and *Mien1* (bottom) (mean±95% confidence intervals of three technical repeats for each sample) in spleens and tumour samples from *BlgCre-NeuKI* mice and in wild type and *NeuKI* organoid controls (eight biological replicates each of three technical replicates) by ddPCR. Samples are arranged from lowest to highest amplification from left to right and colour coded to indicate sample type (Black, control; purple, spleen; blue, sarcoma; yellow, ASQC; green, AME; red, AC(NST)). *Indicates analysis carried out on DNA extracted from FFPE material. For some tumours, both snap frozen and FFPE material were tested.

**Figure S4: Copy number changes in *Atad1* and *Pten* in *BlgCre-NeuKI* tumours.** Copy number of *Atad1* (top) and *Pten* (bottom) (mean±95% confidence intervals of three technical repeats for each sample) in spleens and tumour samples from *BlgCre-NeuKI* mice and in wild type and *NeuKI* organoid controls (eight biological replicates each of three technical replicates) by ddPCR. Samples are arranged from lowest to highest amplification from left to right and colour coded to indicate sample type (Black, control; purple, spleen; blue, sarcoma; yellow, ASQC; green, AME; red, AC(NST)). *Indicates analysis carried out on DNA extracted from FFPE material. For some tumours, both snap frozen and FFPE material were tested.

**Figure S5: Non-negative Matrix Factorisation analysis of RNAseq data.** Estimation of the rank by the R package *NMF* [18]. **(A)** Quality measures computed from 50 runs for each value of rank *k* across the expression dataset. **(B)** Consensus matrices ranked 2 – 6 computed from 50 runs for each value of rank *k* across the expression dataset. The top ranked matrix is shown in **Figure 6A**.

**Figure S6: Relapse Free Survival (RFS) of HER2 amplified and non-amplified breast cancer stratified by canonical WNT target genes.** Data from KM plotter for unselected breast cancers, HER2 amplified and HER2 non-amplified breast cancers stratified by expression of *WIF1* **(A)**, *TCF7* **(B)**, *AXIN2* **(C)** and *MYC* **(D)** using ‘auto select best cut-off’ mode.

**Supplementary Tables**

**Table S1: Full details of tumour cohort animals included in the study.** The database number of the animal, the block number of any sample taken for pathology, age, parity status, number of tumours, location and comments are provided. For a small number of cases histology blocks were missing.

**Table S2: Probes for qRT-PCR, digital droplet PCR and genotyping.**

**Table S3: Full statistics matrix showing statistical differences between tumour cohorts.** Statistics for survival curves (**Figure1A** and **1C**) are provided using both the Log Rank Mantel-Cox test and the Gehan-Breslow-Wilcoxon test. Statistics for differences between tumour numbers per animal **(Figure 1D)** used the Mann-Whitney test.

**Table S4: Full descriptions of tumours which underwent detailed histological examination.** Tumours were assessed as described previously [7,8]. All *K14Cre* and *BlgCre* tumours which were available underwent detailed examination. Only a selection of the *MMTV-NeuNT* tumours, chosen at random, underwent detailed examination. Scoring of staining for IHC markers (K14, K18, ERα, PR, p63) was by estimating percentages of positive cells within the tumour. Negative = no staining; 1 = <10% of cells positive; 2 = 10 – 50% cells positive; 3 = 50 – 75% cells positive; 4 = >75% cells positive.

**Table S5: Details of samples used for exome sequencing and RNAseq.** Sample numbers allocated by the sequencing facilities, tumour numbers, parity status, tumour phenotype and the amplification status of the locus by qPCR are provided, sorted by both tumour number and sequencing number for convenience.

**Table S6: Detailed results of exome sequencing.** Coding mutations identified are listed in the first sheet and the full Mutect and VEP outputs for each tumour are provided. Tumour IDs 11 to 32 refer to the IDs on Table S5 (B67-0011 to B67-0032).

**Table S7: Summarised exome sequencing data.**

**Table S8: Detailed results of CNV-by-exome analysis.**

**Table S9: Summarised CNV-by-exome analysis.** Gains/losses in analysed tumours with an amplification log ratio >4.00 or a loss log ratio <-2.50. Genes in the predicted amplified/lost segment are indicated, together with a summary of their expression levels from the RNAseq data, in order to conclude whether or not the predicted genomic change has potential as a driver event.

**Table S10: ddPCR analysis of *NeoR*, *Neu*, *Erbb2*, *Grb7*, *Mien1*, *Pten* and *Atad1*.** Data for each locus are provided as a mean and 95% confidence intervals (‘Error max’ and ‘Error min’ based on triplicate technical replicates of each tumour and eight biological replicates, each of three technical replicates, for the wild type and *NeuKI* organoids) for the control organoid samples and the tumours. A summary sheet for the *NeoR – Neu – Erbb2* locus is also provided, and a sheet showing the concordance of results across different methods. The majority of analyses were carried out on snap frozen material, however for some samples only DNA extracted from FFPE sections was available. For a subset of samples, the analysis was carried out on both.

**Table S11: Normalised RPKM values for tumours analysed by RNAseq.**

**Table S12: Significantly differentially expressed genes in highly amplified and non/low amplified tumours.** Relative expression levels for all genes are shown as well as genes with a significant (FDR P value <0.05 and a fold change >2) expression difference comparing tumours with a highly amplified with a non/low amplified *Neu* allele. Genes significantly upregulated in highly amplified tumour are, by definition, significantly downregulated in non/low amplified tumours, and vice versa.

**Table S13: Functional annotation and functional annotation clustering of genes differentially expressed in highly amplified and non/low amplified tumours.** Clustering by DAVID v6.8 using GO Bioprocess and KEGG pathway annotation. Genes of Interest (GoIs) associated with the GO/KEGG terms GO:0016055~Wnt signalling pathway, GO:0060070~canonical Wnt signalling pathway, GO:0090263~positive regulation of canonical Wnt signalling pathway, GO:0030216~keratinocyte differentiation, GO:0031424~keratinization, mmu04310:Wnt signalling pathway and mmu04012:ErbB signalling pathway are shown on a separate tab. Log2 RPKM values (from **Table S11**) for the GoIs in the individual tumours are also shown.

**Table S14: Genomic locations of differentially expressed genes.** Genomic locations by chromosome number and locus start site from Mouse Genome Informatics database (JAX).
