## Supplementary figures and images for "Reproductive history determines *ErbB2* locus amplification, WNT signalling and tumour phenotype in a murine breast cancer model"

### Supplementary Figure 1

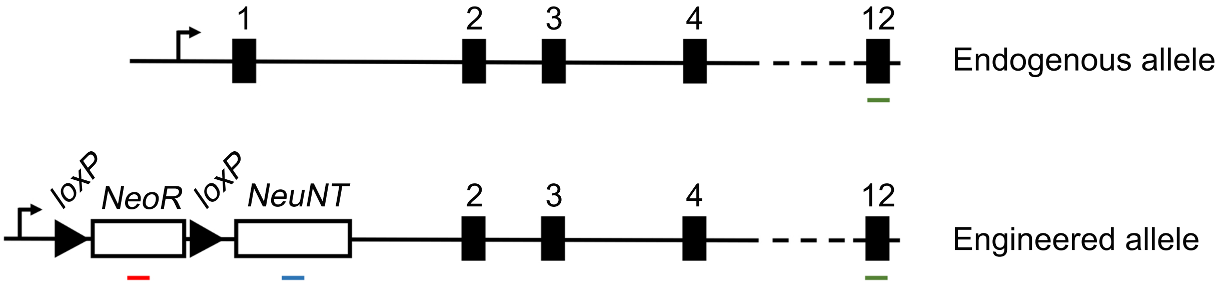

### Supplementary Figure 2

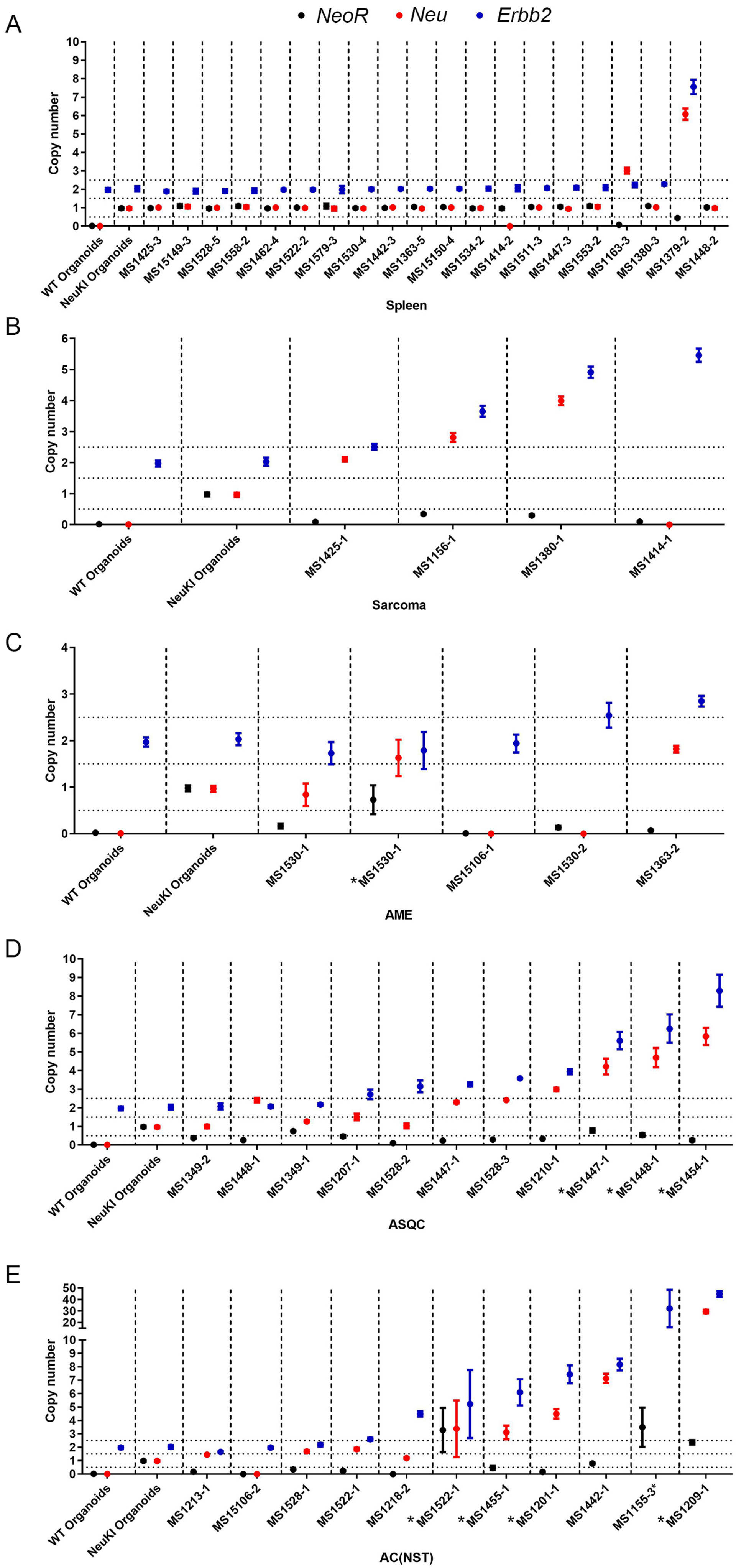

### Supplementary Figure 3

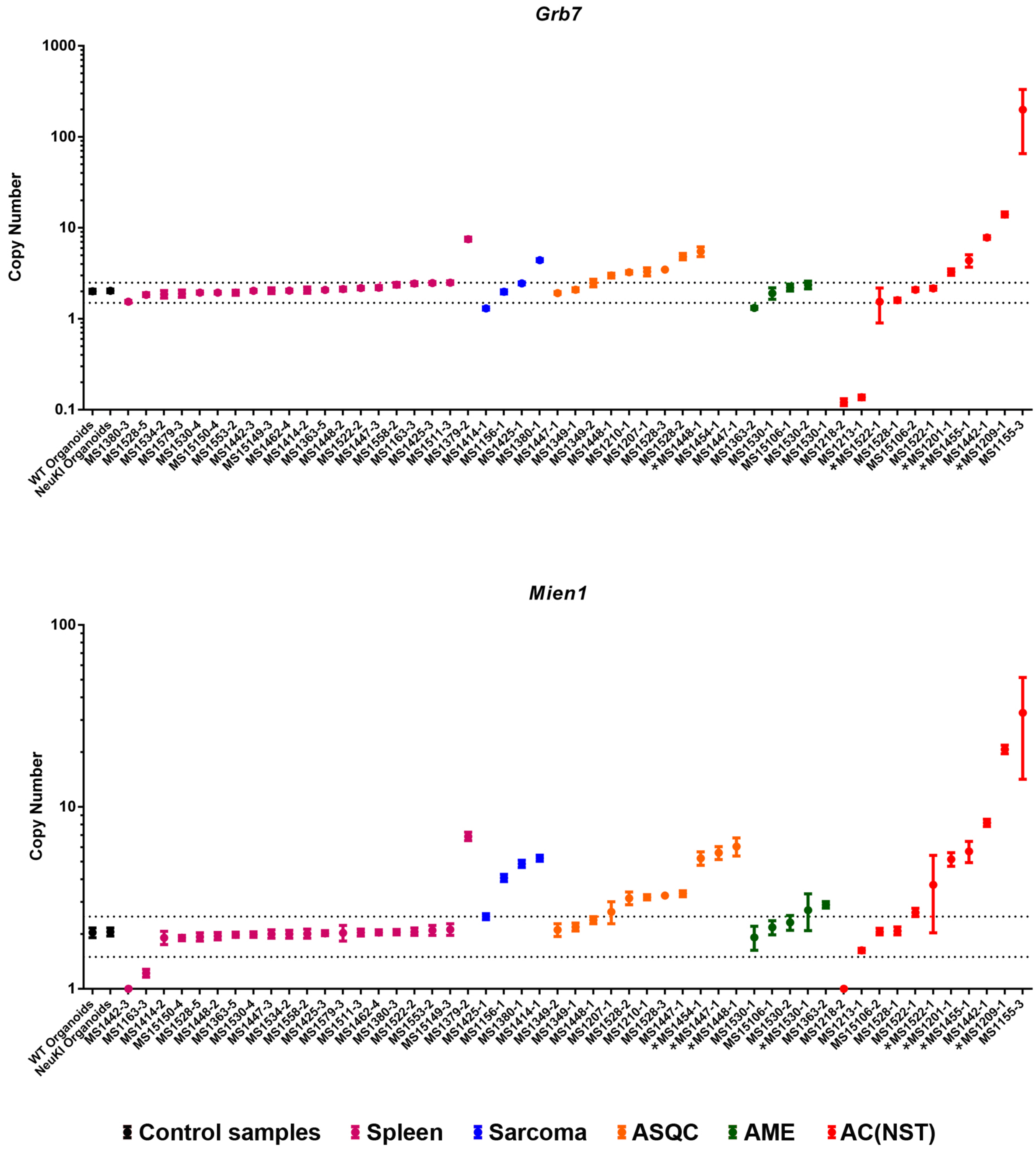

### Supplementary Figure 4

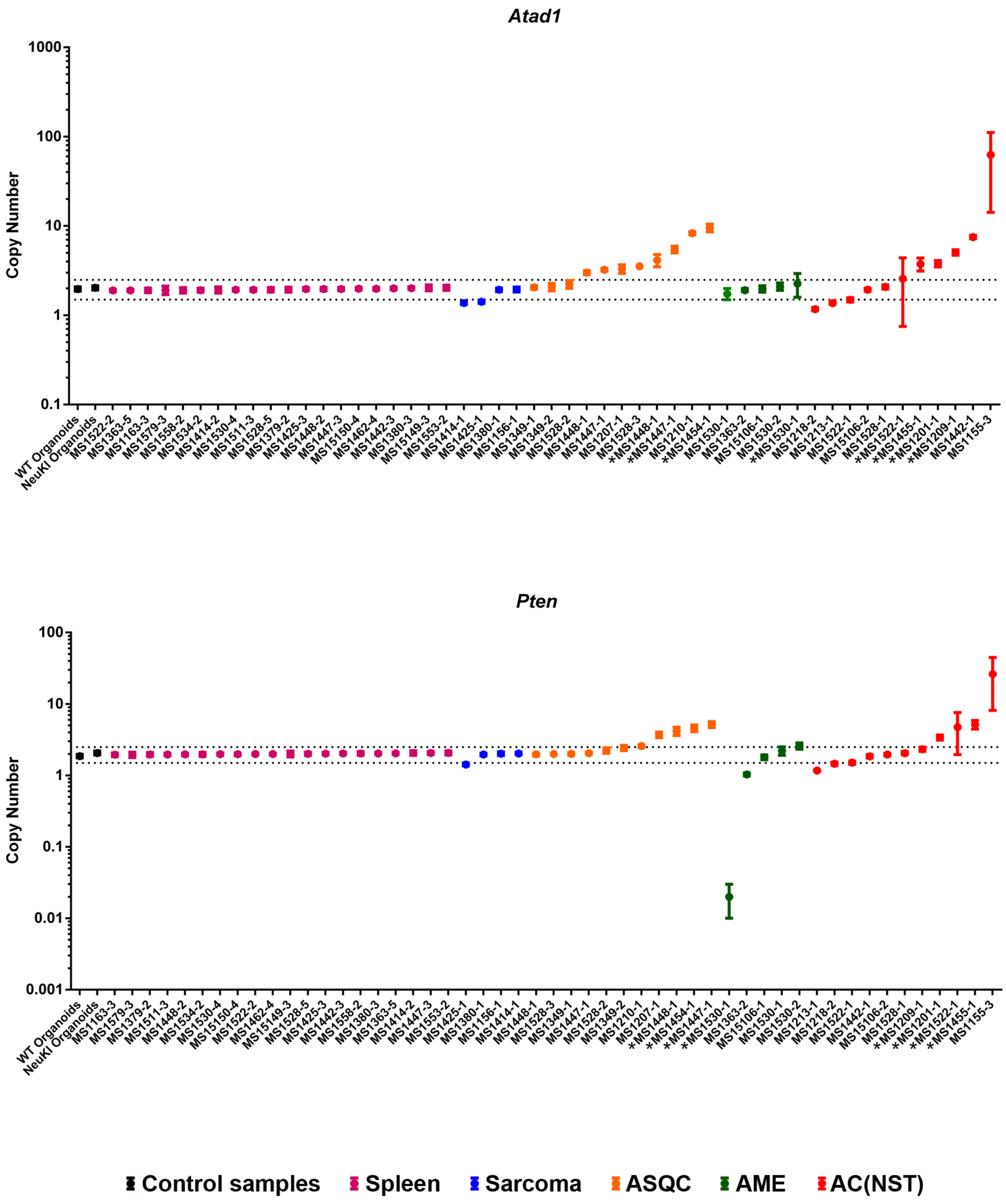

### Supplementary Figure 5

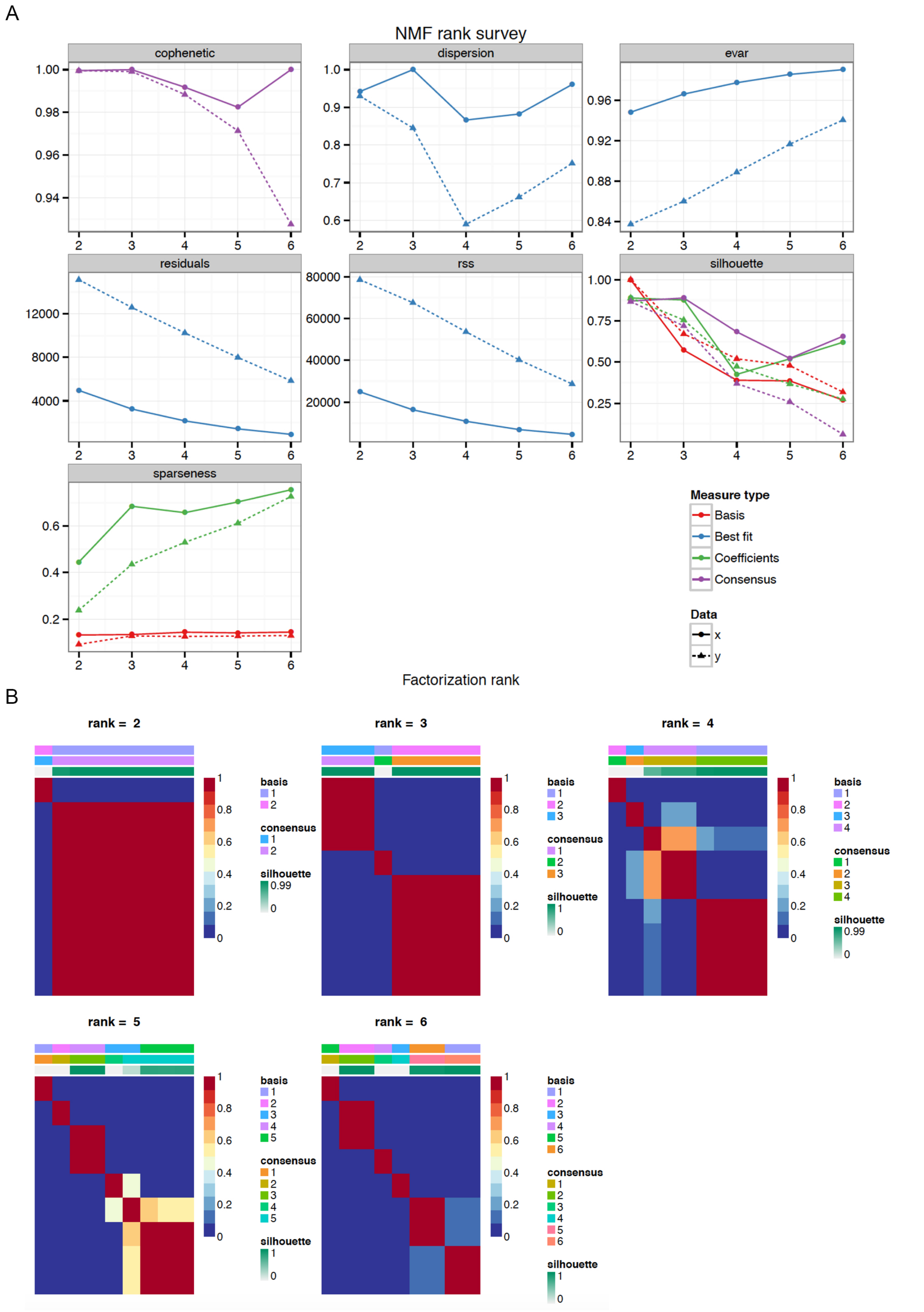

### Supplementary Figure 6

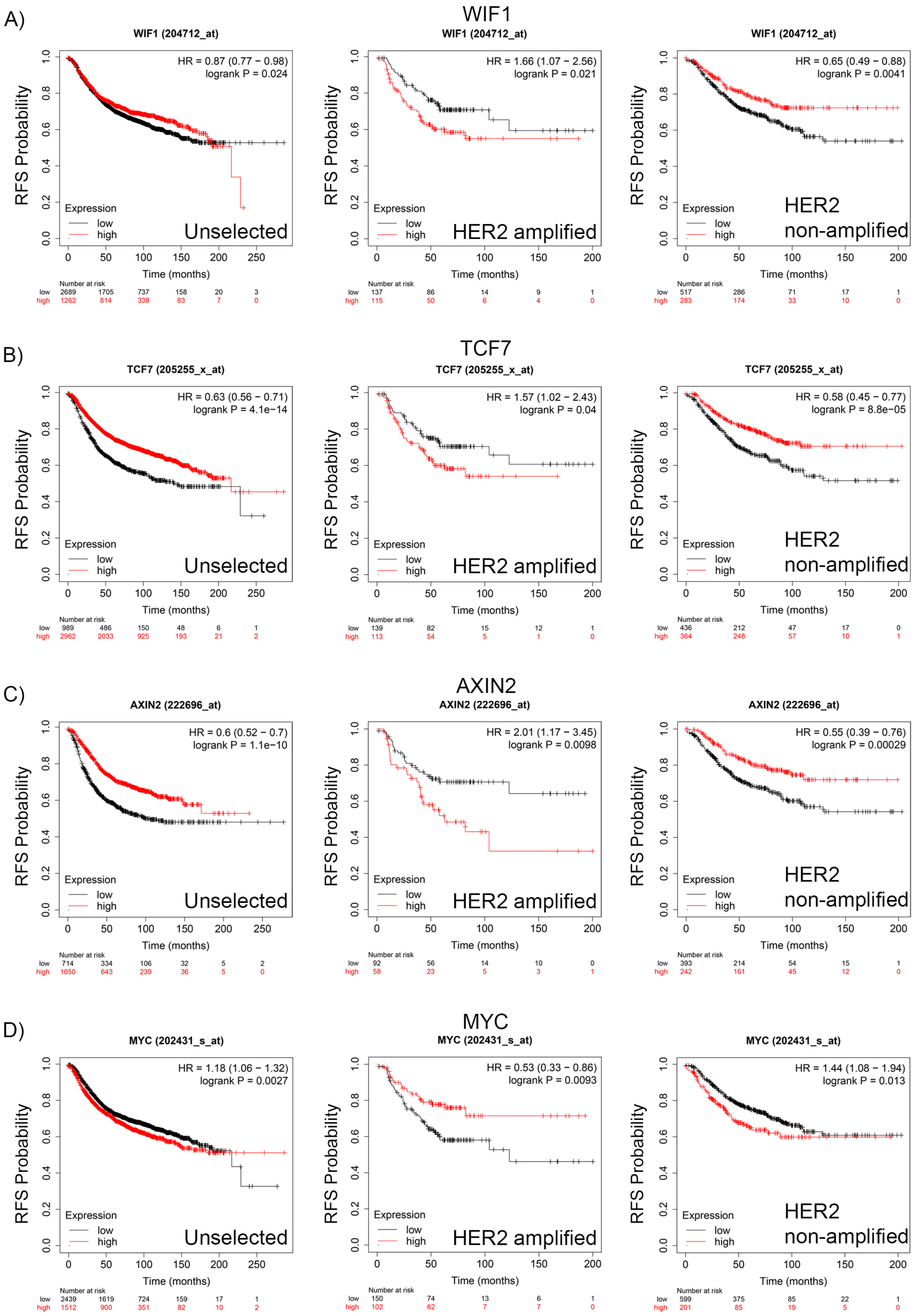
